# Mechanisms of perisynaptic astrocyte depolarization in response to neuronal activity

**DOI:** 10.1101/2024.06.05.597669

**Authors:** Ryo J. Nakatani, Erik De Schutter

## Abstract

Recent studies show that astrocytic depolarization can be induced at the periphery of cortical somatosensory astrocytes, proposed to be the contact sites between neurons and astrocytes. However, specific mechanisms causing astrocytic depolarization have yet to be confirmed due to limitations in experimental techniques. Here, we constructed a computational whole-cell astrocyte model to assess which channels were responsible for astrocyte depolarization. Our simulations show that, unlike depolarization by bath application of potassium, local depolarization by potassium uptake and glutamate transporters required very large spillover and high-frequency stimulation. On the contrary, the model reproduced experimentally observed depolarizations by activating N-methyl-d-aspartate receptor (NMDAR) or γ-aminobutyric acid A receptor (GABA_***A***_R), on the astrocyte. Our models suggest two mechanisms for astrocyte depolarization, either by neurotransmitters or by potassium and glutamate transporters, which substantially alters the spatio-temporal dynamics of the phenomenon. These insights suggest new mechanisms of how astrocytic processes can locally regulate learning and memory.

## Introduction

The electrophysiological properties of brain cells are the foundation of information transfer within the brain. For the past century, neurons have been thought to be the only cell type with these dynamic electrical properties. As active changes of membrane potentials are how neurons propagate information, other non-neuronal cells, otherwise known as glia, were considered irrelevant to the brain’s information processing.

Astrocytes, which account for more than half of glial cells in the human brain, have generally shown passive responses to classical electrophysiological techniques. Such studies have explored various conditions that alter astrocyte membrane potentials, such as changing extracellular potassium concentrations or genetic knock-out (KO) of inwardly rectifying potassium 4.1 (Kir 4.1) channels. Such manipulations result in an elevated resting membrane potential [1, 2]. These experiments have established that astrocytes act like ‘potassium electrodes’, reflecting changes in extracellular potassium. However, most of these studies use bath conditions, which influence whole-cell astrocyte membrane potentials. Local manipulations such as potassium puffs result in minimal observable changes with somatic recordings, unless utilizing excessive concentrations [3, 4]. Therefore, observations have often neglected responses at peripheral locations in the astrocyte, due to technical limitations in electrophysiology.

Recent reports utilizing fluorescent voltage indicators show dynamic changes in membrane potentials caused by neural stimulation that occur only at the peripheral parts of an astrocyte [5]. The depolarizations recorded in these cortical somatosensory astrocytes were cumulative with 100 Hz stimulation and could reach 20 mV after 10 electrical stimuli to ascending excitatory axons. The depolarization sites observed in this paper are suggested to be in the vicinity of where astrocytes and synapses make contact, called perisynaptic astrocytic processes (PAPs). The depolarization is thought to be caused by membrane potential changes at PAPs responding to extracellular potassium elevation induced by action potentials [5, 6].

Recent evidence shows that PAPs are where the astrocyte can modulate neuro-transmission, through gliotransmission or changes in homeostatic functionality [7–9]. Interestingly, the depolarization of astrocytes can contribute to the slowing of gluta-mate uptake by the astrocytic transporters, resulting in spillover [5, 10–12]. Therefore, depolarizations of PAPs have been hypothesized to strongly affect synaptic responses, potentially affecting brain computation.

This raises the possibility of an astrocyte-initiated active response to neuronal activity within the cortex. However, it is still unclear whether these potassium-mediated depolarizations are feasible under physiological neuronal activity, as the observed depolarization would require at least 10 mM of extracellular potassium [5]. As potassium can diffuse to nearby synapses or be taken up by postsynaptic potassium-chloride co-transporter 2 (KCC2) channels, maintaining such a high potassium concentration is thought to be difficult [13] and contradicted by recent reports of only ≈ 1 mM increases caused by neural activity in vivo [14]. The specific mechanisms causing PAP depolarization remain largely unknown. Only major astrocyte channels such as Kir 4.1 and excitatory amino-acid transporter (EAATs), and the *γ*-aminobutyric acid A receptor (GABA_*A*_R) have been observed to contribute to extracellular potassium, although in whole-cell and non-local conditions [5, 15–17]. This lack of experimental data, along with experimental limitations preventing direct observation of PAPs, requires the use of computational approaches.

In this study, we constructed an empirical conductance-based NEURON model with a realistic morphology in order to capture both PAP and whole-cell astrocyte properties. Using this model, we explored possible mechanisms to alter astrocyte membrane potentials dependent on neuronal activity, trying to match experimentally observed changes. Experiments estimate a peak depolarization of 20 mV in peripheral sub-compartments upon ascending-axon stimulation in the somatosensory cortex [5], and so we tried to match this value. First, we recreate how astrocytes react to potassium in classical manipulations, such as bath application, to establish our model. We next examine how these responses change with local potassium application, with a focus on single synapse level activity. As our results indicate difficulty in depolarization by potassium alone, we next examine how neurotransmitter transporters on the astrocyte contribute to astrocyte depolarization. Our findings show that this is a feasible mechanism of depolarization, but only under high-frequency stimulation followed by large amplitudes of spillover and potassium accumulation. We finally explore other mechanisms that could depolarize the astrocyte with minimal and more realistic changes of global extracellular conditions. Our model suggests astrocytic neurotransmitter receptors to be a viable candidate, causing strong astrocytic depolarization at perisynaptic microdomains. Our model provides a crucial hypothesis on the active regulation of PAP depolarization and suggests new ways in which astrocytes can modulate synaptic function.

## Results

### Potassium induces depolarization in astrocytes dependent on global extracellular changes of potassium

We constructed an astrocyte potassium whole-cell model to examine the interaction between potassium and the membrane potentials of an astrocyte. The constructed conductance-based model contained the major contributors of potassium, as well as other channels relevant to astrocytic depolarization using a previously reported whole-cell morphology [18] (Fig. 1 A,C). This morphology contained astrocyte nanoscopic structures that recreate characteristics of PAPs, such as the higher input resistance when compared to the soma (Supplementary Fig. 1 A). We did not consider gap junctions in our model as we were more interested in local changes at PAPs that contacted synapses rather than the periphery attached to other astrocytes. The model we constructed was specifically designed with a focus on membrane potential and extracellular potassium, which required channel models to be dynamically responsive to changes in potassium reversal potential. Therefore, previously established astrocytic channel models with built-in membrane potential sinks were excluded from our model [19]. To validate our model, we tested the astrocyte response to patch clamp protocols, which showed passive responses as reported in classic literature [20] (Fig. 1 B). The model exhibited a hyperpolarized resting membrane potential (RMP) between −80 ~ −85 mV, which is another defining characteristic of astrocytes [21].

**Fig. 1:**
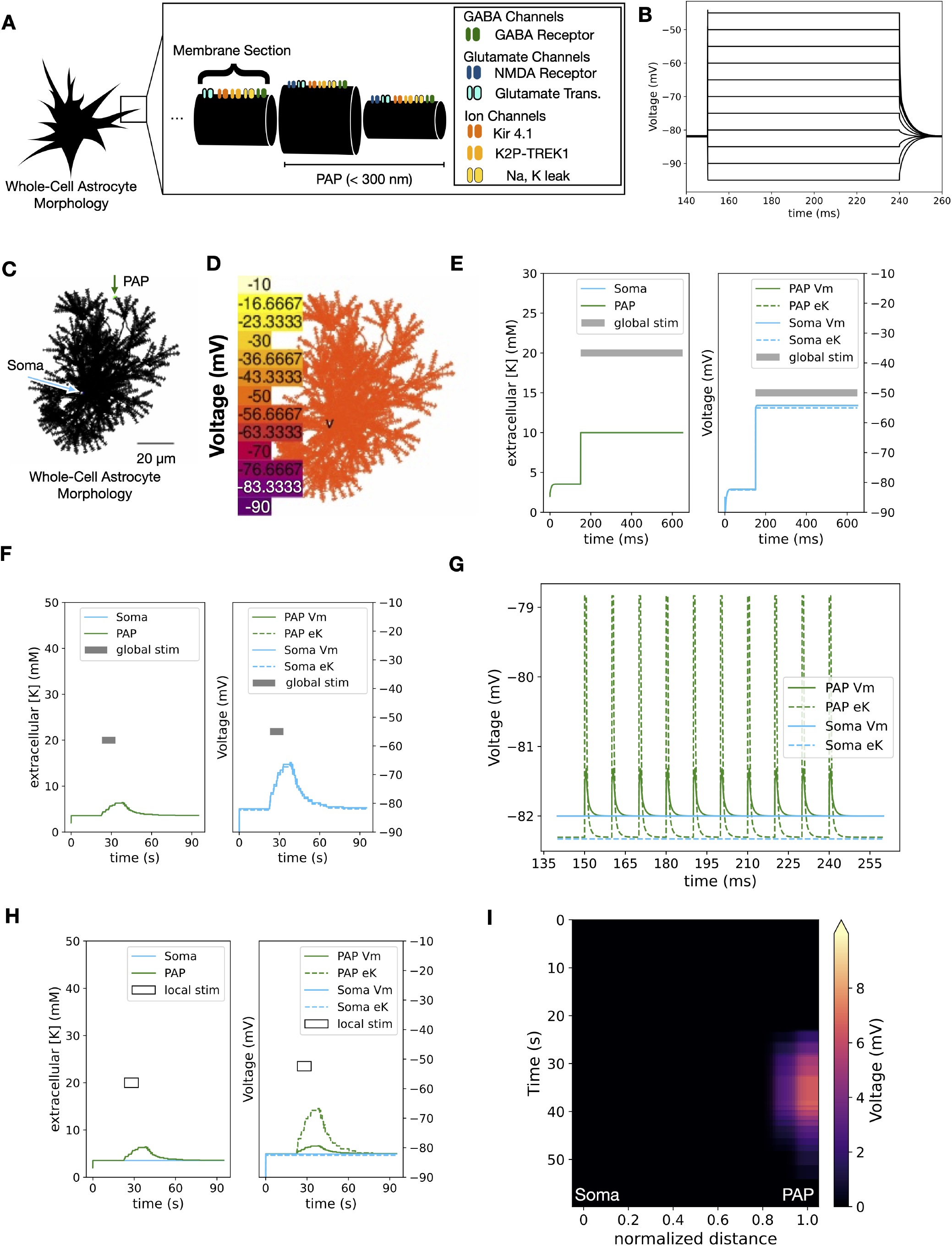
The potassium-driven whole-cell astrocyte model (A)Cartoon depicting our conductance-based model. The astrocyte morphology is that of a previously constructed model [18], with new equations for Kir 4.1 and K2P-TREK1 channels and for glutamate transporters (GluTs). Only portions of the model up to 300 nm in length and diameter less than 0.5 µm were considered a PAP candidate. These sections were randomly selected, excluding the primary branch or soma, unless stated otherwise. As the synaptic receptor distribution was unknown, only PAP sections had synaptic receptors. For all other channels, all distributions were uniform to match densities defined for the PAP. All sections have a sodium leak current to maintain RMP. Each cylinder in the cartoon represents a cable section of the astrocyte model. See also Supplementary Fig. 5 and supplementary text 1 ~ 7 for justification of channel models. (B)The soma voltage response to various voltage clamps in our model. Voltage commands are −95 ~ −45 mV with5 mV increments, which show typical passive features of the astrocyte. (C)A typical representation of the PAP and soma within the whole-cell morphology of the astrocyte previously described in Savtchenko et al. [18]. The scale bar shows 20 µm. (D)Peak depolarization visualized under bath application of potassium. The color bar indicates the membrane potential of each compartment and corresponds to the color scale within the figure. The depolarization is induced with 482 Kir channels on the PAP, and an extracellular potassium concentration of 10 mM. (E)Changes in extracellular potassium and membrane potential for both soma and PAP during the bath application protocol (panel D) over time. After bath application of 10 mM extracellular potassium, the membrane potential instantaneously changes, following the change in potassium Nernst potential. The resulting depolarization reaches ~ 20 mV. (F)Changes in extracellular potassium and membrane potential for both soma and PAP reflecting the *in vivo*protocol. The experimental *in vivo* protocol [2] results in a slow change of extracellular potassium over the timecourse of seconds. Similarly to panel E the membrane potential follows changes in Nernst potential. (G)Currents during the synaptic stimulation of simple APs (sAP; 0.5 mM changes of potassium held for 0.5 ms). Potassium is not accumulated during 100 Hz single synapse simulations, and the membrane potential change is less than 1 mV. (H)Changes in extracellular potassium and membrane potential in changes isolated at the PAP reflecting the *in vivo* protocol. Unlike panel F, the changes in membrane potential are below the Nernst potential for the PAP. (I)Voltage attenuation during the *in vivo* protocol. The distance between the soma and PAP is normalized on the x-axis, and the changes in these sections are plotted over time. Membrane potential changes are rapidly attenuated, creating a large voltage leak in the PAP. Any voltage changes in the PAP did not affect the soma.

In order to examine the relationship between extracellular potassium and astrocyte membrane potential, we first examine how our model interacts to bath application of extracellular potassium. Previous literature shows that the membrane potential of astrocytes follows that of the potassium Nernst reversal potential [6]. As expected, our model also depolarizes following the Nernst reversal potential with bath application of 10 mM of potassium (Fig. 1 D,E).

As our model was able to recreate astrocyte depolarizations caused by global extracellular changes, we questioned whether large depolarizations were possible with physiological brain stimuli. By applying simplified action potentials (sAP; 0.5 mM increase of extracellular potassium for 0.5 ms) for 10 pulses at 100 Hz with a normal density of Kir 4.1 channels, we sought to investigate how astrocytes respond to an active single synapse. Therefore, changes in extracellular potassium due to stimulation were isolated in the PAP. The stimulation protocol was chosen to match the stimulus protocol from previous experimental work [5]. The potassium change caused by a single action potential (AP) was estimated using values from previous computational models of potassium in the synaptic cleft [13, 19]. Potassium contributions from postsynaptic neuronal *α*-amino-3-hydroxy-5-methyl-4-isoxazolepropionic acid receptor (AMPAR) and N-methyl-D-aspartate receptor (NMDAR) activity were regarded as negligible for distances of astrocytes from the postsynaptic density (PSD) [22]. As the repolarization process of neurons typically lasts ≈ 0.5 ms, a period of 0.5 ms was chosen.

The sAP stimulation resulted in minimal depolarizations that did not match experimental values, as there is no potassium build-up within the extracellular space at the PAP (Fig. 1 G). The lack of accumulation of extracellular potassium in our model was due to the quick removal of potassium, mainly done via astrocytic Kir 4.1 (Supplementary Fig. 1 C). In this protocol we selected a PAP with the largest response to 16 mM change of extracellular potassium, but the response was still within two standard deviations from the mean response of randomly selected PAPs (mean: −65 mV, *σ*: 2.8 mV; Supplementary Fig. 1 D). AP responses also varied from location to location, although the qualitative characteristic of not being able to follow the Nernst reversal potential remained the same (Supplementary Fig. 1 E).

In order to further investigate how differences in global versus local application affect depolarization, we next recreated experimentally observed long-term *in vivo* changes of extracellular potassium within our model [2]. In doing so, we sought to examine how these potassium changes would trigger different astrocyte responses in both global and local contexts. For global application, *in vivo* extracellular potassium changes also induced depolarization within our model, which followed direct changes to the reversal potential, similar to our bath application protocol (Fig. 1 F vs. E). However, upon confining the potassium changes to PAPs, we observed a substantial decrease in depolarization amplitude in comparison to their globally modulated counterparts (Fig. 1 H vs. F). Furthermore, our results show that the lack of potassium build-up was not the sole reason for the low amplitude, which was mainly caused by the difficulty in causing isolated depolarization. Our model indicates strong attenuation (space constant 4.56 µm) with no voltage changes observed at the soma (Fig. 1 I), suggesting that local depolarization is strongly diminished by the voltage ‘leak’. These results suggest that potassium and major astrocytic potassium channels alone are insufficient for PAP depolarization in conditions using physiological stimulation of single synapses.

### Potassium induced focal depolarizations require large amplitudes of extracellular potassium

As our previous results highlighted the difficulty of depolarizing astrocytes with local physiological stimuli, we next theoretically explored various AP scenarios to explore possibilities that could locally depolarize the astrocyte. First, we examined the relationship between the density of Kir 4.1 channels and the extracellular potassium stimulus amplitude, with a range covering from little Kir 4.1 or no extracellular potassium changes to unphysiologically high densities and potassium amplitudes. The results showed a strong potassium dependence, with higher potassium concentrations than calculated from the Nernst potential and Kir 4.1 densities required for significant depolarizations (Fig. 2 A). In order to match experimental values of peak PAP depolarization [5] with our sAP stimuli we found that ≈ 20 mM was necessary to reach the experimental values, higher than the initially proposed 10 mM (Fig. 2 A,F).

**Fig. 2:**
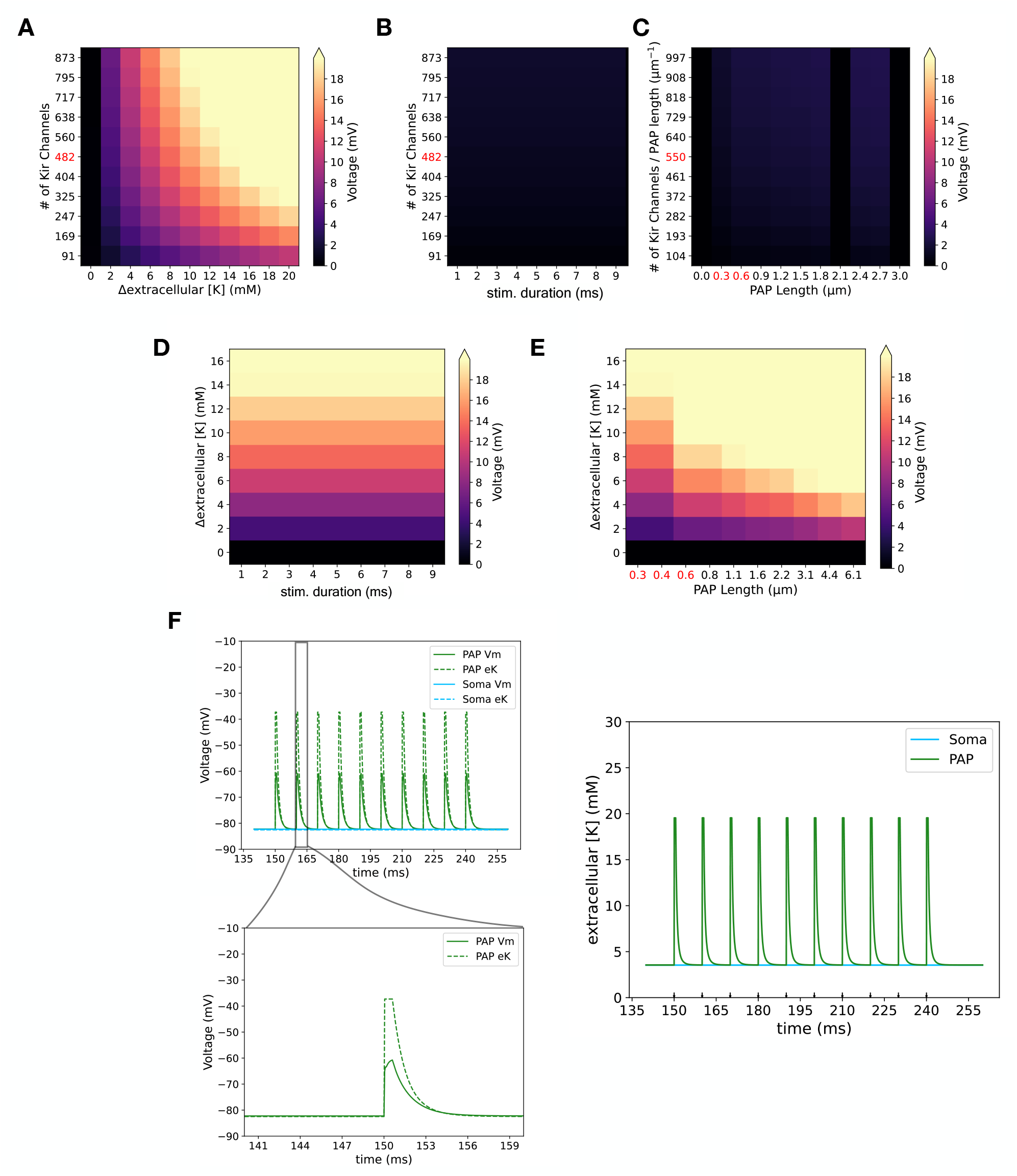
Exploration of various potassium stimuli scenarios **(A)** Heat maps showing maximum depolarization comparing each count of Kir 4.1 channels to different amplitudes of extracellular potassium stimuli. The peak depolarization was recorded at the PAP with a single stimulus lasting 0.5 ms. The colors indicate the peak membrane potential for each condition, corresponding to the color scale within the figure. The red text label indicates the conditions within the standard deviations measured from experimental results (Kir 4.1; 3.7 × 10^8^ *±* 10^8^ channels per cm^2^ [50]). The PAP length used was 0.3 µm, which was the standard protocol. **(B**,**C)** The peak depolarization for different stimulus conditions comparing differences in Kir 4.1 channel numbers to various B) stimulation duration and C) PAP length. As the altered PAP length changes the number of recruited Kir 4.1 channels (panel C), channel numbers were normalized to per µm of PAP length. The potassium amplitudes were 0.5 mM for all conditions. **(D**,**E)** The peak depolarization for different stimulus conditions comparing differences in potassium amplitudes to various D) stimulation duration and E) PAP length. The Kir 4.1 channel count was 482 for all conditions. **(F)** Depolarization of astrocyte by 20 mV using amplitude modified sAP protocol. The sAP had a modified potassium stimulus of 16 mM (total of 19.5 mM in the extracellular space) with a duration of 0.5 ms. Left) Membrane potential changes over time in response to the modified sAP, with an inset showing the changes in membrane potential at the single stimulus level. Right) Changes in extracellular potassium during the protocol over time. Simulations were done with 482 Kir 4.1 channels on the PAP.

Next, we considered alteration in the duration of the 0.5 mM potassium stimuli, as well as the size of the potassium affected area. These changes reflect different AP scenarios, such as long-tailed APs or closely clustered synapses, which result in potassium release at a larger spatial scale. We examined a stimulus duration until 10 ms, which was equivalent to the inter-stimulus interval, as well as PAP lengths about a sixth of the length of the primary branch for protoplasmic astrocytes [23]. The results showed that the duration of the potassium stimuli contributed minimally

(Fig. 2 B). Similarly, changes in the potassium-affected area also did not substantially alter peak depolarization amplitudes (Fig. 2 C). Our results suggest that the amplitude of potassium is the largest driver of depolarization, with astrocyte-affected length contributing as well (Fig. 2 D,E). In order to reach experimentally observed values, large amplitudes of potassium (≈ 20 mM) or PAP affected lengths larger than typical PAPs were necessary for considerable depolarization (Fig. 2 D-F).

### GluT driven depolarization of astrocytes requires spillover of glutamate

Our previous results show that stimuli with extracellular potassium changes alone will not substantially depolarize the astrocyte unless with unphysiologically large amplitudes of extracellular potassium changes. As experimental results observe more isolated depolarization, this suggests that there are other mechanisms that contribute to astrocyte depolarization. These studies have shown that glutamate transporters, or GluTs (GLT-1), contribute to astrocytic depolarization, with an estimate of about 20% of the peak depolarization (approximately 3 ~ 4 mV) [5]. In order to test this hypothesis, we examined depolarization of our model with only GluT. For locally confined changes of extracellular glutamate at the single synapse level, we saw that astrocytic depolarization was not affected even with an increase of GluT density by 5 times the standard deviation of estimated physiological values (Fig. 3 A).

**Fig. 3:**
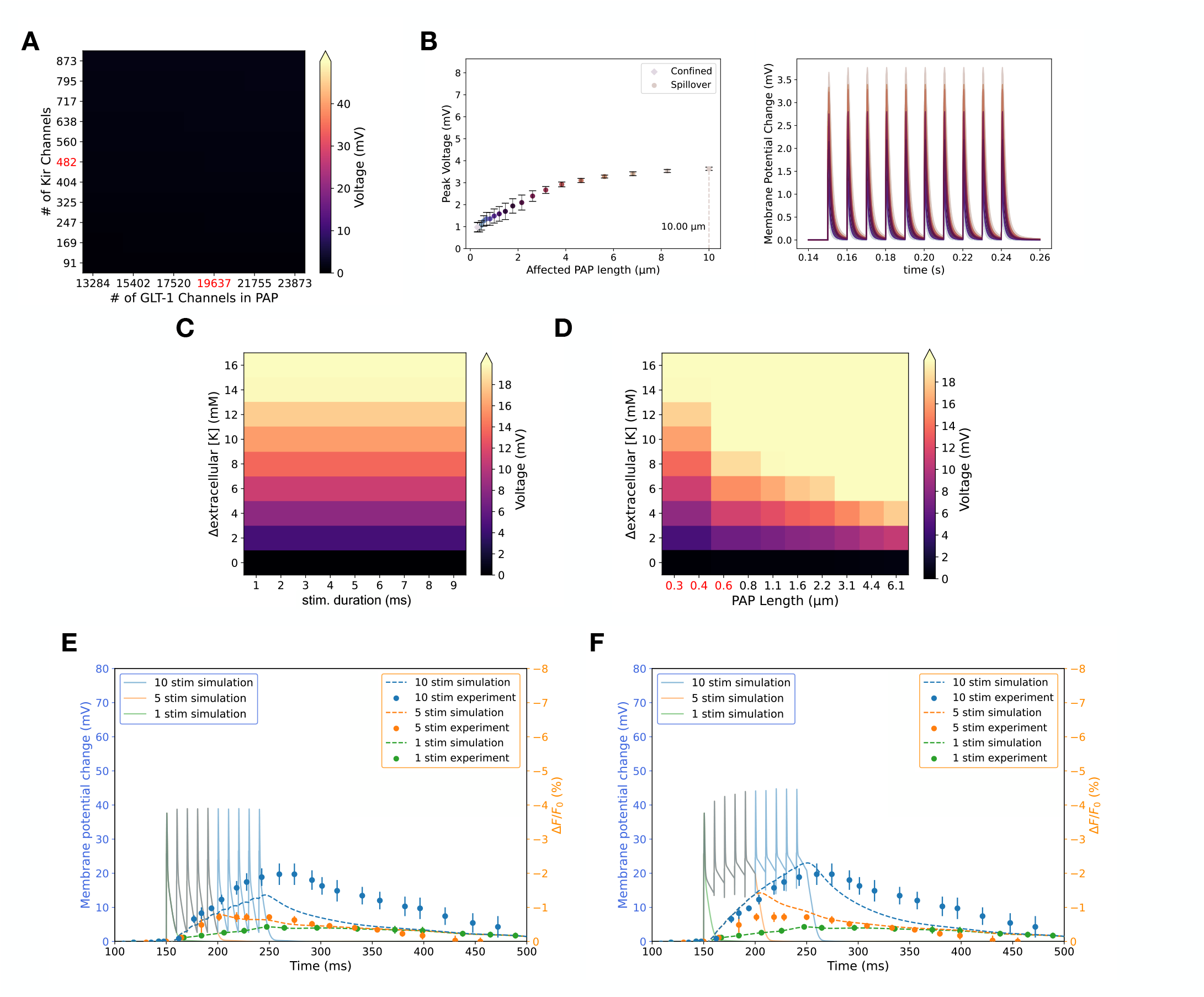
Glutamate transporter (GLT-1) driven depolarization **(A)** Heat maps showing maximum depolarization comparing differences in counts of Kir 4.1 channels and glutamate transporters. The heat-map shows peak depolarization recorded at the PAP with single stimuli of 0.5 ms. The colors indicate the membrane potential for each condition, corresponding to the color scale within the figure. The red text labels indicate the conditions within the standard deviations (Kir 4.1; listed previously, GLT-1; 1.4 × 10^12^ *±* 8.1 × 10^10^ per cm^2^ [60]). **(B)** Left) Plots of peak depolarization against PAP lengths. The diamond shown in the plot and legend indicates the physiological conditions of the affected PAP length. Longer lengths than the physiological conditions are all spillover conditions, which are marked with a circle. n=10 for randomly selected PAP locations for each data point. The PAP length for the largest observed depolarization is marked on the x-axis with its PAP length value (10 µm). Right) Currents of each individual condition for plots on the right. Each trace color corresponds to that in the right plot. **(C**,**D)** The peak depolarization for different stimulus conditions comparing the effects of potassium amplitudes with the GluT potassium model. C) Stimulation duration and D) PAP length were altered for each respective plot. For panel C the PAP length was maintained at 0.3 µm. The Kir 4.1 channel count was 482, and 19637 for GLT-1 channels for all conditions. **(E**,**F)** Model fitting to experimental results observing peripheral depolarization in astrocytes [5]. The left y-axis (blue) shows membrane potential changes within our model, and the right y-axis (orange) shows changes in the fluorescent voltage indicator for both simulated and experimental values. The slow accumulation of voltage by the single stimulus protocol (green data points) was considered an artifact, as it could not be recreated by other fluorescent voltage indicators. Therefore, the artifact was fitted with a polynomial spline and then added to all simulated fluorescent traces. E) Conditions only altering peak amplitudes of potassium and neurotransmitters, as well as affected PAP lengths, result in a poor fit (SSE: 8.42). F) Conditions also altering decay dynamics of potassium and neurotransmitters result in a better fit to the results (SSE: 4.61). Glutamate decay time constants was changed to 10 ms from 5.8 ms, and potassium decay constant was altered to 40 ms from 4 ms.

We next investigated how glutamate spillover would affect depolarization, as spillover could recruit more GluTs. We used a simplified model of local spillover of glutamate by increasing the affected PAP length, and examined peak depolarizations for various affected lengths in our conductance-based model. As a consequence, we did not consider 3-dimensional aspects of extracellular diffusion. Upon increasing the length of the glutamate-affected PAP, we find that for lengths above 2 µm there are depolarizations of over 2 mV (Fig. 3 B). Our model could achieve experimental values, but only at lengths over 4 µm, analogous to when there is a strong ‘spillover’ of glutamate. This suggests that depolarization of astrocytes via GluTs can only be triggered with strong repeated stimuli that have largely synchronized release/spillover of glutamate.

By combining both potassium-mediated depolarization with GluT depolarization, we next estimated the amplitude of depolarization that could be achieved. Our model shows that GluTs contribute minimally, showing a larger contribution from changes in extracellular potassium (Fig. 3 C,D). We next fit our model to previously conducted Arclight experiments [5], which have a measured calibration curve converting fluorescent changes to membrane potential changes. By implementing ordinary differential equations that match the time dynamics of Arclight within our simulations, we acquired pseudo-fluorescence changes reflecting our model’s membrane potential responses.

As a result, we show that without significant spillover and consistent accumulation (i.e., slowed decay of neurotransmitters) of neurotransmitters and potassium, experimental traces could not be matched (Fig. 3 E,F). In the latter case, only the amplitudes but not the decay could be matched to the experimental data (Fig. 3 F). This suggests that these mechanisms can only contribute to depolarization under high-frequency stimulation with near-pathological extracellular conditions.

### Neurotransmitter mediated depolarization

In many pathological cases where neurons are hyperactive, astrocytes have been observed to depolarize along with brain-wide increases in extracellular glutamate and potassium [24]. Our previous results suggest that astrocyte depolarization is not only induced in pathological conditions but also in conditions that can trigger local ‘spillover’. However, we questioned whether astrocytic depolarization could only be triggered by stimuli inducing spillover, such as high-frequency stimulation. This difference would suggest a more active role in astrocytic depolarization in comparison to being a local by-product of spillover. Previous results show that sub-compartments of astrocyte activity are coupled to neuronal activity, suggesting that astrocytes can respond to neurotransmitters [25–27]. One such receptor is the GABA_*A*_R, which is expressed on astrocytes and is known to depolarize astrocytes [15, 16]. The contradiction to typical neuronal GABA_*A*_R roles is due to the different intracellular chloride concentration of astrocytes, leading to depolarizing GABA_*A*_R currents [16, 28].

We therefore next examined how GABA_*A*_Rs could hypothetically contribute to local depolarization of astrocytes, and questioned what depolarization amplitudes could be observed in astrocyte sub-compartments that are parallel to synapses. We implemented a classic two-state neuronal inhibitory model of GABA_*A*_Rs, with a minor alteration to the chloride reversal potential to fit the astrocyte profile. We examined the peak depolarization in the PAP by adding 1.5 mM of GABA after the potassium stimulus to recreate neurotransmitter release [29]. Using the same 10 pulse 100 Hz stimulus of sAPs, we observe depolarizations of approximately 20 mV, with physiological densities of astrocytic GABA_*A*_R (Fig. 4 A,B). Although the depolarization was sustained for longer than that of Kir 4.1 only, voltage attenuation maintained the isolation of PAP voltage dynamics (Fig. 4 C). Only upon changing the PAP location to somewhere closer to the soma (*<* 2 µm), could we see changes in soma membrane potential (Supplementary Fig. 2 B). GABA_*A*_Rs also produced depolarization that did not require large potassium amplitudes or affected PAP lengths, which were key factors for astrocyte depolarization in our previous results. On the other hand, during the depolarization of the PAP by GABA, the membrane potential rose higher than the potassium reversal potential, resulting in an increase of extracellular potassium (Fig. 4 D). Global application of GABA also resulted in minimal somatic changes, which matched classic experimental studies (Supplementary Fig. 2 A). These results suggest that GABA_*A*_R may also contribute to astrocytic depolarization in brain regions with abundant inhibitory neurons.

**Fig. 4:**
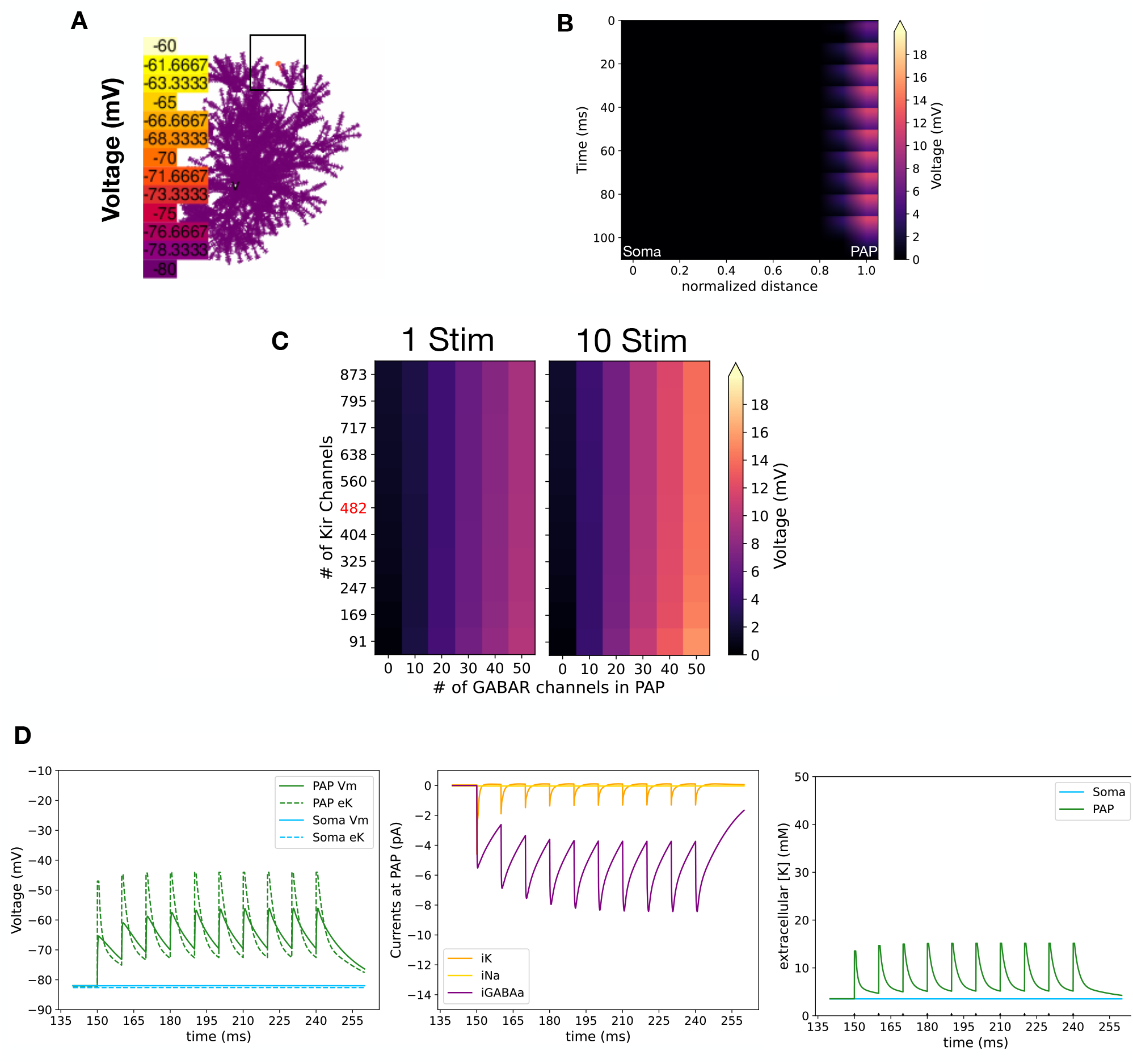
GABA_*A*_R driven depolarization **(A)** Peak depolarization visualized in our whole-cell astrocyte model. There are 80 GABA_*A*_Rs and 482 Kir channels on the PAP. The sAP stimulation protocol with 1.5 mM of GABA for 10 pulses at 100 Hz was used for depolarizing the astrocyte. **(B)** The voltage attenuation along the branch from soma to PAP. Distance is normalized to the distance of the PAP from the soma. **(C)** Heat maps showing maximum depolarization comparing each count of Kir 4.1 channels and GABA_*A*_R. The peak depolarization recorded at the PAP was compared for both single and 10 stimuli of 0.5 ms. All stimuli were followed by 1.5 mM increase of GABA. The colors indicate the membrane potential for each condition, corresponding to the color scale within the figure. The red text labels indicate the conditions within the standard deviations. There were no absolute estimates for the number of GABA_*A*_Rs on astrocytes available. **(D)** Left) Voltage change along with reversal potential (eK), Middle) currents in PAPs, Right) extracellular potassium concentration within the tip of the PAP section. No glutamate transporters were activated during this protocol. Parameters for these plots were chosen to match those of panel A.

Although GABA-mediated depolarization suggests astrocyte membrane responses to inhibitory neurons, it was logical to ask whether this response was selective to inhibitory synapses. Recent studies have shown behavioral changes induced by knock-down (KD) of cortical somatosensory astrocyte NMDARs, suggesting that these channels are functionally expressed [27], with direct implications for astrocyte calcium and function. Therefore, to observe the effects of NMDAR, we implemented a previously reported NMDAR model [30] with low density on the PAP. We then examined the peak depolarization in the PAP by using the same stimulation protocol as GABA_*A*_R depolarization, except for replacing GABA with glutamate. As astrocytic NMDARs have various differences to neuronal NMDARs, such as less susceptibility to Mg^+^, we tuned the parameters of the NMDAR model to recreate depolarization amplitudes observed in experimental results [5] (Fig. 5 A,D). As a result, we found that NMDAR-mediated depolarization replicates our GABA_*A*_R results in terms of isolation of the stimuli (Fig. 5 B). This isolation was maintained regardless of the number and location of PAPs (excluding soma or primary branch of the astrocyte) simultaneously triggered, with activation of 10 randomly selected PAPs resulting in a *<* 1 mV change at the soma (Supplementary Fig. 3 B). However, it is important to note a few key differences when comparing to GABA_*A*_R-mediated depolarization (Fig. 5 A-C). Firstly, NMDAR depolarization seemed to have larger accumulatory effects, which are similar to experimental results [5], as NMDARs have longer channel open times than GABA_*A*_Rs (Fig. 5 B,C). The NMDAR depolarization, therefore, matches the threshold like response of NMDAR spikes which have a strong on/off response to a single stimulus, which is not the case for the GABA_*A*_R response. Secondly, the change in channel open times also affected the decay time-course of depolarization and extracellular potassium (Fig. 5 C). Lastly, our model shows that depolarization via NMDAR can be done with a smaller number of channels compared to that of GABA_*A*_R (Fig. 5 A). We therefore suggest that NMDAR channels that are expressed at a much lower density than Kir 4.1 channels or GABA_*A*_Rs can still depolarize the astrocyte to the same amplitudes as GABA_*A*_Rs. The NMDAR model also matched fairly well with the experimental results, suggesting that PAP depolarization amplitudes could be achieved without strong spillover or accumulation of neurotransmitters (Fig. 5 D). Additionally, we observed that the weak Mg^+^ block produces NMDA spike-like features to the depolarization with minimal changes to astrocyte membrane potential for the first and second stimuli (Fig. 5 B,D). This effect was also observed when altering synaptic frequency and pulse numbers, as small numbers and slow frequencies resulted in minimal changes (Supplementary Fig. 3 A).

**Fig. 5:**
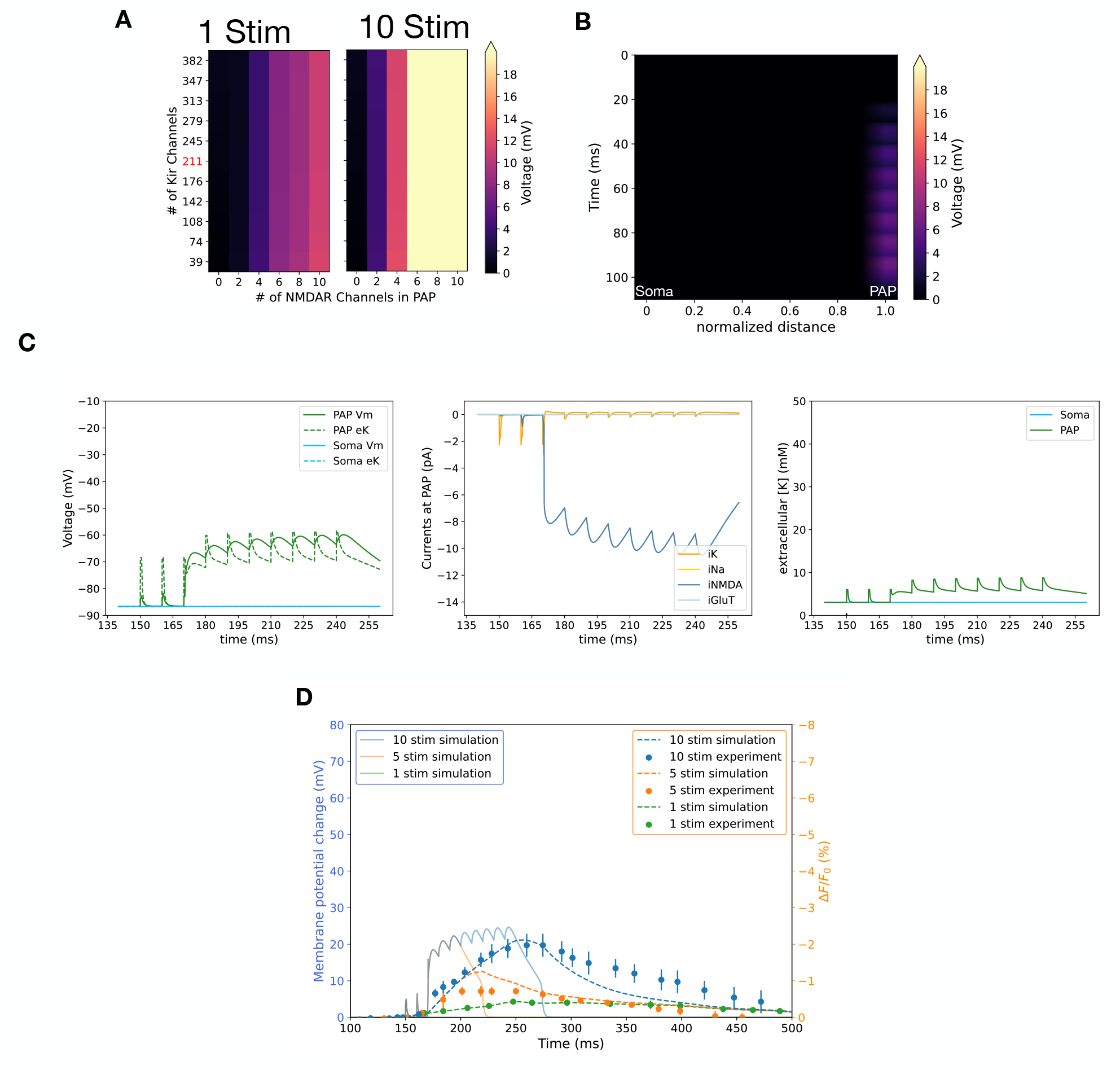
NMDAR driven depolarization **(A)** Heat maps showing typical depolarization response of PAPs with various counts of Kir 4.1 channels and NMDAR. The peak depolarization recorded at the PAP was compared for both single and 10 stimuli of 0.5 ms. All stimuli were followed by 1 mM increase of glutamate. The colors indicate the membrane potential for each condition, corresponding to the color scale within the figure. The red text labels indicate the conditions within the standard deviations. There were no absolute estimates for the number of NMDARs on astrocytes available. The PAP location differs from previous experiments in this protocol. **(B)** The voltage attenuation along the branch from soma to PAP. Distance was normalized to the distance between the PAP far end and the soma. **(C)** Left) Voltage change along with reversal potential (eK), Middle) currents, Right) extracellular potassium concentration within the tip of the PAP section and soma. The parameters used in this plot match the fit results of panel D. **(D)** Model fitting to experimental results observing peripheral depolarization in astrocytes [5]. The left y-axis (blue) shows membrane potential changes within our model, and the right y-axis (orange) shows changes in fluorescent voltage indicator for both simulated and experimental values. The slow accumulation of voltage by the single stimulus protocol (green data points) was considered an artifact. Therefore, the artifact was fitted with a polynomial spline and then added to all simulated fluorescent traces. Conditions with no spillover using glutamate produce fluorescent traces that follow the experimental results (Sum of squared error; SSE: 3.20).

### Astrocyte channel composition modulates depolarization amplitude

So far, our results suggest that potential mechanisms of astrocyte depolarization can be classified into two large categories. The first category is that of potassium/GluT-mediated depolarization, which depends on high-frequency neuronal activity and spillover of neurotransmitters and potassium. On the other hand, neurotransmitter-mediated depolarization shows depolarization amplitudes of the same level with minimal spillover. The differences between the two suggest that astrocyte depolarization responses could be heterogeneous depending on the channel composition of the PAP. Therefore, we examined how differences in channel composition could alter depolarization amplitude.

First, in order to examine the influence of all channels, we examined how astrocyte membrane potential responses under high-frequency stimuli differ in various knock-out (KO) conditions. This comparison showed that the NMDARs strongly control the depolarization peak whenever they are present, with depolarization enhanced by glutamate spillover conditions (Fig. 6 A). On the other hand, GABA_*A*_R mediated depolarization was diminished under spillover conditions, due to the lack of GluT activation as well as a stronger activation of Kir 4.1 channels (Fig. 6 A). This was confirmed with Kir 4.1 over-expression in our model, which lowers depolarization amplitudes for NMDAR-mediated depolarization (Supplementary Fig. 4). Interestingly, these results match experiments that have tested Kir 4.1 overexpression (OE) conditions [5].

**Fig. 6:**
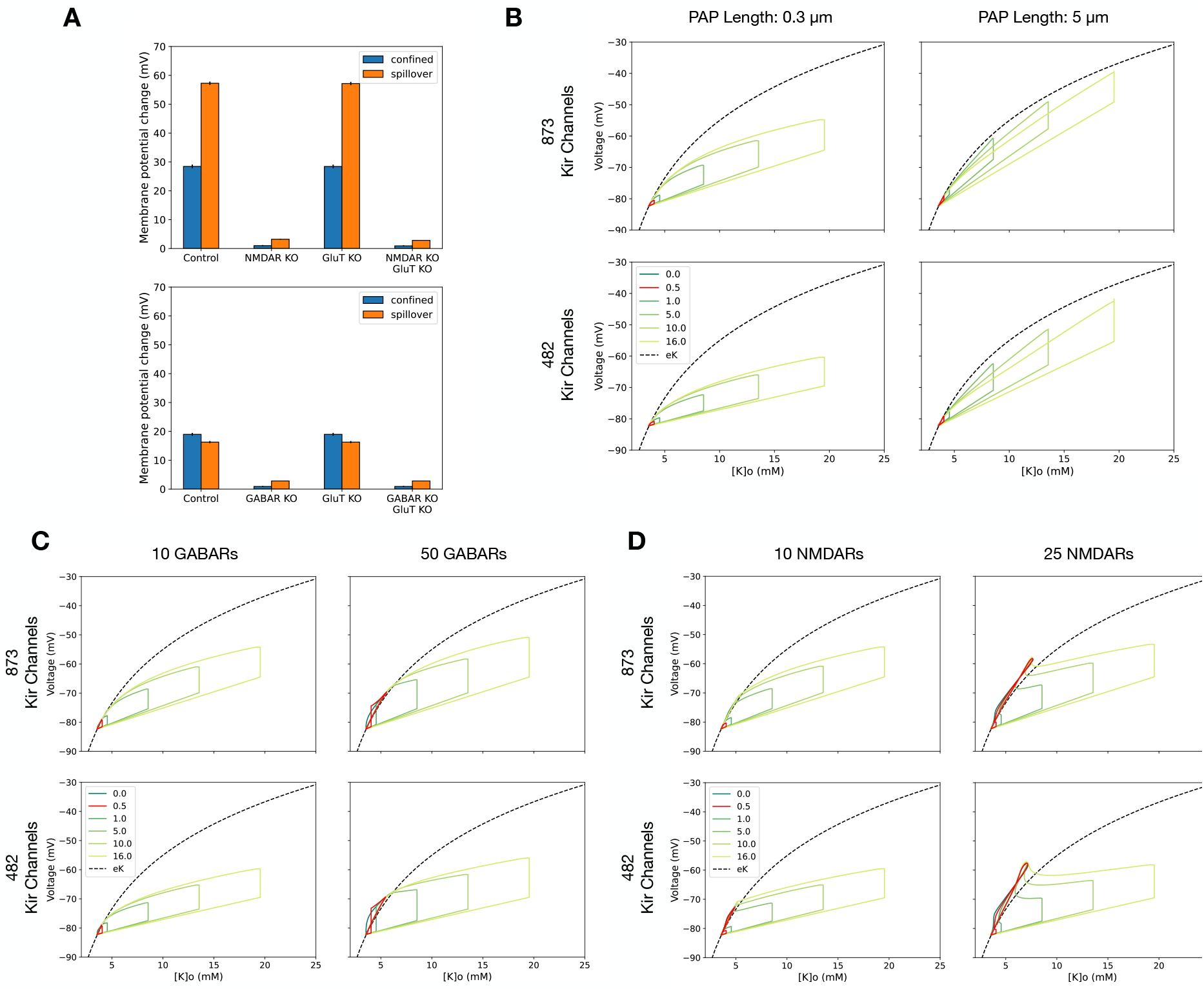
A comparison of various channel densities for PAP depolarization **(A)** Comparison of peak depolarization for knockout (KO) conditions of synaptic receptor KO, GluT KO, and GluT/synaptic receptor double KO. All experiments were conducted for GABA_*A*_R and NMDAR, respectively. All conditions were stimulated with the sAP protocol, followed by a neurotransmitter release of 1 mM glutamate or 1.5 mM GABA. Control conditions were done with PAP lengths of 0.3 µm, while spillover conditions had affected PAP lengths of 5 µm. The y-axis shows the peak depolarization for each condition, with spillover causing a larger depolarization for all KO conditions in glutamate stimulated conditions (upper panel). Differences between confined and spillover were statistically tested for all conditions, with results shown in Supplementary Table 8. **(B-D)** Phase planes of extracellular potassium ([K]o) vs. voltage for respective combination of B) PAP lengths, C) GABA_*A*_R counts, and D) NMDAR counts. Traces for phases with different amplitudes of extracellular potassium stimuli are color coded and labeled in the legend. Each panel contains four figures with the corresponding conditions labeled for each column and row. All phase planes show the model responses to a single stimulus and compare phase portraits of extracellular potassium versus membrane potential. The control condition for relative extracellular potassium increase has been colored in red. The reversal potential calculated from voltage and extracellular potassium concentration is plotted as a broken line in black.

To further analyze how each channel contributes to depolarization dynamics, we examined the phase plane of extracellular potassium against membrane potential for each of our models (i.e. Kir4.1/GluT, GABAR, NMDAR). This revealed how channel densities, as well as PAP lengths, affected the depolarization dynamics of PAPs (Fig. 6 B-D). We revealed that generally more activated synaptic receptors and GluT induce larger depolarization amplitudes, while large Kir 4.1 densities create stronger sinks to the reversal potential. Differences in potassium stimulus amplitude also affected depolarization, although not as much as altering the number of synaptic receptors. In regards to the Kir4.1/GluT model, we saw that depolarization amplitudes were much smaller compared to other models, with changes in PAP length and activated GluT channels not affecting depolarization as much as changes in synaptic receptor numbers (Fig. 6 B). The primary changes caused by increasing NMDAR/GABA_*A*_R were large increases in potassium concentration and membrane potential, specifically seen as a loop structure in the upper left of the phase plane (Fig. 6 C,D; left vs. right). Therefore, our results suggested that NMDARs contribute most efficiently to depolarization, with Kir 4.1 channel and NMDAR balance strongly determining the depolarization dynamics.

## Discussion

This study focused on exploring possible mechanisms responsible for peripheral depolarization of cortical astrocytes using a new empirical conductance-based mathematical model. The construction of a new conductance-based model was necessary as we wanted to examine astrocyte channel numbers with unitary conductances. This also allowed for a more detailed investigation of the Kir 4.1 channel and potassium currents, unlike other previous Kir 4.1 models [19] that included a built-in leak factor.

This study also examined popular hypotheses regarding changes in extracellular potassium as well as glutamate transporter currents, and compared them to responses mediated by synaptic receptors. We believe that this study showcases how each hypothesis can play a role in contributing to depolarization, with some mechanisms highly dependent on high-frequency stimulation and extracellular build-up, while others could be evoked with spontaneous neuronal activity. Our model showed that although Kir 4.1 alone can depolarize the PAP, it required ≈ 20 mM of potassium for physiological Kir 4.1 densities, which was a very high external potassium concentration that was difficult to achieve using single synaptic events (Fig. 1). Moreover, potassium is cleared rapidly and is not expected to accumulate during the 10 ms interval between stimuli. It was also observed that extracellular potassium changes alone in isolated compartments could not increase the membrane potential to calculated Nernst potential values because there was a constant voltage leak. We further examined this model with the implementation of GluTs, which also did not contribute significantly and required strong accumulation of neurotransmitters to contribute (Fig. 3). However, depolarization by synaptic receptors significantly contributed to the isolation of membrane potential changes. Interestingly, the stimulus count-dependent facilitation was largely different depending on which synaptic receptors were activated. GABA_*A*_R had less accumulation of depolarization during stimuli, while NMDAR activation was a slow process that is well known to build up with repeated stimulation [31].

We believe that our model highlights two novel insights into astrocytic electrophysiology. Our first point is that the dynamics for whole-cell and local periphery are vastly different, with conventional notions like the Nernst potential being insufficient when considering dynamics at the nanoscopic level (Fig. 1). The second is that astrocytes may use synaptic receptors to alter membrane potentials, which would dynamically alter astrocyte responses to neuronal stimuli (Fig. 4, 5). These insights together imply that astrocytes may have dynamic neuronal responses that are invisible to conventional electrophysiological techniques. We believe that although the study by Armbruster et al.[5] was the first step to identify these hidden astrocyte responses, a more thorough examination discriminating whole cell and local differences is necessary. Our model indicates that in order to achieve similar results, utilizing potassium mechanisms, requires high-frequency stimulation and strong accumulation in extra-cellular space, which may not be a commonly occurring phenomenon in physiological brain conditions. Indeed, in vivo recordings of mice somatosensory cortex L2/3 during running show ≈ 1 mM increase in extracellular potassium, highlighting the difficulty to achieve confined large amplitudes of potassium in physiological settings [14]. It also requires attention that results from Armbruster et al. [5] omit the possibilities of astrocytic synaptic receptors because they apply 2-amino-5-phosphonopentanoic acid

(APV) for all experiments and only stimulate excitatory axons. Their experiments used to observe specific channel contributions were also only analyzed with whole-cell averages, neglecting possible distribution differences in nanoscopic domains. Any receptor that would be clustered in peripheral subcompartments of the astrocyte would be neglected, as the membrane area is much smaller than that of the whole cell. This leaves the possibility for further examination of local astrocytic response to neuronal activity, using techniques with better spatial resolution such as expansion microscopy or stimulated emission depletion (STED) imaging. The apparent absence of effect of APV on the experimentally observed ROI depolarizations [5] may also be due to differences in astrocyte NMDAR subunit composition, as certain NMDAR subunits are less susceptible to APV [32], which may also account for reports on astrocyte NMDA-induced, APV-insensitive currents [33, 34]. The low and heterogeneous expression of astrocytic NMDAR has been discussed previously [35, 36].

Our model suggests that both excitatory and inhibitory neurotransmitters result in similar effects in regard to astrocyte membrane potentials. This suggests that astrocytes can respond to neuronal activity without regard to neuron type. Some studies suggest astrocytes can integrate neuronal activity, for both inhibitory and excitatory synapses, where calcium transients have been observed for both responses [37, 38]. From our results, we speculate that astrocytic ionotropic synaptic receptors can trigger similar responses for both inhibitory and excitatory neurotransmitters that may contribute to calcium transients. We observed a complex interaction scheme between Kir 4.1 and synaptic receptors, where synaptic receptors produced a depolarization that triggered an efflux of potassium. Initially, in the steady state with no excessive extracellular potassium, the Kir 4.1 channel produces a weak outward current that balances leak currents (Supplementary Fig. 7 A). However, the reversal potential for potassium (eK) is altered upon an increase in extracellular potassium produced by neuronal activity (sAP). Kir 4.1 produces a strong instantaneous potassium-dependent inward current, pushing the membrane potential to that of the new eK (Supplementary Fig. 7 B). As the application of neurotransmitters follows the potassium stimulus, this triggers the synaptic receptor to activate, pushing the membrane potential over the eK and reverting Kir 4.1 back to an outward current that increases extracellular potassium (Supplementary Fig. 7 C,D). This depolarization and accumulation of extracellular potassium continues while the synaptic receptor current is stronger than the Kir 4.1 outward current (Supplementary Fig. 7 D). Eventually, the time course of depolarization follows the interplay between the synaptic receptor inward current and Kir 4.1 outward current (Supplementary Fig. 7 D,E) until the next stimulus triggers an additional extracellular potassium increase (Supplementary Fig. 7 B). In regions of clustered spines, we suggest that both excitatory and inhibitory synapses would depolarize the astrocyte membrane as well as trigger transients in extracellular potassium, which would significantly change astrocyte functions in that local spatial domain. At the same time, we believe this could trigger changes in presynaptic release probabilities for spines located near the astrocyte. In this sense, astrocyte function and neuronal computation could be spatially coupled to specific dendrites that integrate input, with strong isolation from other regions within the astrocyte.

PAP depolarization may also strongly alter astrocytic function in information processing [5]. This could be achieved by the voltage dependence of GLT-1, as well as contributing to calcium dynamics by altering Na/Ca exchanger functionality [39]. However, the extent to which the mechanism affects brain computation is unknown. The prevailing assumption is that astrocyte computation is done through increases of intracellular calcium [40, 41]. Although at first glance, PAP depolarization may seem to drive calcium influx, we believe depolarization does not directly contribute to activating voltage-dependent calcium channels (VDCC). This is because VDCCs on astrocytes (L-type, T-type, N-type, R-type) have high voltage thresholds that are difficult to achieve with ranges of depolarization shown experimentally [5] and in the model [42]. Previous experimental results match this description, where astrocyte L-type VDCC-mediated influx required bath application of 75 mM extracellular potassium, expected to cause large membrane depolarization [1]. When considering the possibility of dynamic astrocyte synaptic responses, we believe it would be possible for the astrocyte to tailor its response based on the specific synapse it is in contact with. For example, the number of NMDARs in each PAP may be actively regulated by protein trafficking, where specific PAPs are tailored to have more NMDARs. Protein trafficking in astrocytes is not well understood, but certain proteins like glutamate transporters and aquaporins (AQP4) have been observed to be trafficked via anterograde vesicle transport [43, 44].

An alternative way astrocytes may affect brain computation is through the manipulation of extracellular potassium, observed during depolarization of our synaptic receptor astrocyte model (Supplementary Fig. 7). These changes in extracellular potassium are thought to strongly alter neuronal excitability [14], and thus, astrocyte-mediated potassium regulation may actively alter synaptic transmission in local and neighboring synapses. Note that, because of the rectification properties of the Kir 4.1 channel, the potassium outflow will stop increasing if the PAP is depolarized even more, limiting the predicted potassium elevation.

Our current findings suggest that PAP depolarization can be triggered by both high-frequency stimulation and synaptic receptors. With the whole-cell conductance-based model, we showed K^+^/GluT mediated mechanisms required strong spillover and high-frequency stimulation, while synaptic receptor-mediated depolarization could occur with spatial isolation. For K^+^/GluT mediated depolarization, we show that depolarization requires potassium amplitudes that are much higher than initially anticipated by previous reports [5]. As the impact of PAP depolarization on synaptic plasticity is still unknown, further investigation into the properties of glutamate diffusion at the nanoscopic spatial scale is required.

## Materials and Methods

### Model design

The objective of the study was to construct a realistic whole-cell astrocyte *in silico* model, capable of recreating the peripheral depolarization phenomenon. Therefore, the model required dynamic updates of the Nernst reversal potential reflecting changes in extracellular potassium. This excluded the use of channel models that had built-in mechanisms for maintaining membrane potential, often used for modeling astrocyte electrophysiology [19]. The morphology used in this model was from Savtchenko et al., which used hippocampal CA1 protoplasmic astrocytes [18]. Morphological differences between protoplasmic hippocampal astrocytes and cortical somatosensory astrocytes are minimal, with high correlation of morphological characteristics between the two [45]. The model morphology is defined in NEURON [46], in the file GeometryAstrocyteCA1.hoc. See also Supplementary Fig. 1 A,B for additional morphological details. All subsequent channel models were defined in NEURON MODL [46]. All parameters used in the model are listed in Supplementary Table 1-7. Any alteration of channel counts was done by altering the single channel conductance linearly. Kir 4.1, GluT, TREK1, leak channels were uniformly distributed for all simulations based on the channel counts defined in the PAP. For GABA_*A*_R and NMDAR, any change is directly reflected in channel numbers solely in the PAP. Custom or redefined channels (Kir 4.1, TREK1) were fitted to experimental data shown in Supplementary Fig. 5. Channels that have the same dynamics as the original model are explicitly cited, and no parameters related to the dynamics were altered, except for the NMDAR receptor, which was fit to experimental results in Armbruster et al. [5]. All referenced MODL files can be found in the neuronMOD file within the uploaded repository.

### Extracellular space

The extracellular space was modeled as a shell around the cellular compartment with an effective thickness of 100 °A. The extracellular diffusion between sections was not considered, as the selected PAP regions were thought to be isolated by neuronal uptake[13]. The difference between basal potassium concentrations of 2.5 mM and the excess amount of potassium was cleared at a time constant of 4 ms. The time constant for the simulation models both the uptake from the extracellular space by various channels on the pre/post-synaptic terminals and the diffusion of potassium away from the astrocyte. These values were fit to maintain physiological extracellular potassium concentrations in the extracellular space.

### Kir 4.1

The Kir 4.1 channel model is adapted from Yim et al. [47] with a [K^+^]_*o*_ dependent square-root law [48].

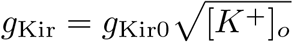

where *g*_Kir0_ is the maximum conductance. Maximum conductance was calculated to fit single-channel recordings and extracellular potassium concentrations of that experiment [49]. Details can be found in the Kir2.mod file and Supplementary text 1.

Estimates for Kir 4.1 channel densities were derived from Bovzic et al. and were about 3.7×10^8^±1×10^8^ channels per cm^2^ [50]. Specific derivations are outlined in Supplementary text 2. Kir 4.1 channel numbers for OE experiments were matched with the observed maximum ratio between control and OE, for experimentally measured currents from Armbruster et al. [5].

### K2P-TREK1 channel

The GHK-dependent rectification-type potassium channel equations were taken from Janjic et al. [51]. The original equations were converted into NEURON and matched the dynamics in the paper. Details can be found in the TWIK.mod file and Supplementary text 3.

### NMDAR

The NMDAR model is adapted from Moradi et al. [30] with a few alterations to fit astrocyte NMDAR recordings. First, the voltage dependence of time constants for the NMDAR model has been refit to that of astrocyte NMDAR recordings by [52]. Second, the magnesium dependence of the model was reduced, as astrocyte NMDAR currents are independent of extracellular magnesium concentrations [52]. Third, the induced glutamate concentration affects the model in a Hill equation-dependent manner, reducing activation when there is a low concentration of released glutamate. Last, the maximum conductance for single channel conductance was calculated from Jahr et al. [53]. NMDAR channel numbers were calculated in a Gaussian-dependent manner for experiments increasing the length of PAPs. This was under the assumption that NMDAR resides mainly at the fine processes [54]. PAPs during spillover experiments contained the most NMDAR at the initial 0.3 µm length and declined with units of 0.3 µm. Details can be found in the SynExp5NMDA.mod file and Supplementary text 4.

### GABA_A_R

The GABA_*A*_R model was a classic two-state synaptic conductance model adapted from Schulz et al. [55] with changes to the reversal potential to fit astrocytic conditions. As GABA_*A*_R reversal potentials are that of chloride [56, 57], we calculated the chloride reversal potential from intra/extracellular concentrations using the Nernst equation. This resulted in a reversal potential of approximately −40 mV. Details can be found in the inh.mod file and Supplementary text 5.

### Glutamate Transporter

The glutamate transporter model (for GLT-1) utilized in this study was taken from that used in Savtchenko et al. [18] and modified from a membrane mechanism to a point process. The model was also changed to affect potassium dynamics and ik, in order to observe how the channel contributed to potassium dynamics of the astrocyte. The channel was observed to have extracellular potassium and voltage-dependent uptake [12] (See also Supplementary Fig. 6). Details can be found in the GluTrans.mod file and Supplementary text 6.

### Leak channels

Leak channels for Na^+^ were modeled as generic conductance-based channels with a predetermined conductance. The specific conductance value was arbitrarily fit, so the resting membrane potential was in the range of −80 ~ −90 mV when Kir channel counts were at a physiological density on the astrocyte PAP. Further explanations of the calculations are detailed in Supplementary text 7.

All leak files are defined as [ion name]leak.mod, where ion names are k or na.

### Ion current to concentration conversion

The ion flux from intracellular to extracellular compartments is calculated with the conversion,

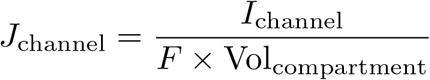

where *J, I* describe the flux and current of any channel, *F* is the Faraday constant and Vol is the volume of the compartment. For potassium and sodium, this is modeled as a NEURONMODL file that follows chapter 9 of the NEURON book [46], modeling Ca^2+^ extracellular concentrations, replacing Ca^2+^ with respective ion properties. All ion conversion files are defined as [ion name]Accum.mod, where ion names are potassium or sodium.

Chloride concentrations in the extracellular space were constant and we did not consider local changes.

### Stimulation Protocols

Before stimulation, the model was subjected to an equilibrium phase where extra-cellular potassium and respective currents achieved a steady state. This equilibrium phase was done for 150 ms when all channel conditions had reached equilibrium. Bath application and *in vivo* stimulation protocols were recreated in the model by clamping extracellular potassium concentrations for appropriate durations. Changes in extracellular potassium for *in vivo* stimulation followed experimental results found in Chever et al. [2]. Either PAP-designated sections or the whole cell was targeted for local and global stimulations, respectively. Plots for PAPs in this paper used seed 1, unless stated, for randomly selecting a location as it was the PAP with the largest response to potassium stimuli (Supplementary Fig. 1 D,E). The selected PAP allowed an easier visual comparison between protocols but was within two standard deviations from the average PAP response. The mean peak response to a 16 mM stimulus of extracellular potassium was −65 mV and the standard deviation (*σ*) was 2.8 mV.

Stimulation protocols using simplified action potentials (sAPs) for depolarizing the astrocyte were done by altering conductances for synaptic receptor models as well as changing extracellular potassium concentrations. The sAP stimuli were triggered at PAP-designated astrocyte sections at 100 Hz to match experimental protocols [5]. Each stimulation was broken down as a 0.5 mM potassium increase that lasts for ms and a 1 mM glutamate or 1.5 mM GABA increase unless stated otherwise. These extracellular potassium increases are confined to the extracellular space of the peripheral astrocyte, which is defined as the PAP. In the case of spillover experiments, this extracellular space is extended, resulting in a longer extracellular space affected by extracellular potassium increases.

### Model fitting to experimental results

The model was fit to experimental results by selecting unconstrained parameters such as synaptic receptor channel numbers, potassium amplitude, and PAP size. Explicitly for GluT-mediated depolarization with forced accumulation, the glutamate decay factor was additionally considered to be a free parameter. In the case of NMDAR, the channel parameters for the magnesium block and open-close time constants were fine-tuned from the original neuronal model, as astrocytic NMDARs have weaker susceptibility to magnesium [52]. All fitted parameters were used for any subsequent simulation to observe the dynamics of the model. Fitting was performed using the Nelder-Mead gradient search algorithm in the scipy package [58] with arbitrary initial values. As the changes in experimental fluorescence for single stimulus protocols were thought to be specific to the used fluorescent voltage indicator Arclight, these traces were considered to be experimental artifacts[5]. This unknown artifact was fit using a polynomial spline and then added to all simulated fluorescence changes.

### Simulations and analyses

All models were implemented in NEURON, where the default astrocyte model was constructed using NEURON HOC version 8.2.0. Specific simulation programming was done by accessing the HOC object in the Python interface using custom code. MPICH [59] was used to parallelize individual simulations, using a custom code for repetitive simulations.

## Acknowledgements

We thank Professor Yukiko Goda for helpful discussions throughout the project and her helpful feedback on the manuscript. We also thank Dr. Jules Lallouette, Dr. Maria Vazquez Pavon for helpful insights on our manuscript.

## Author contributions

R.J.N. and E.D.S conceived the research. R.J.N. constructed the model with assistance from E.D.S.. R.J.N. wrote the original manuscript with suggestions from E.D.S..

## Data and materials availability

The data generated and/or analyzed during the current study are available in the github repository, https://github.com/CNS-OIST/NEURONPAP.

The custom Astrocyte NEURON model created in this study is freely available through modelDB, https://modeldb.science/2019878.

## Supplementary materials

Supplementary Text 1 to 7

Supplementary Fig. 1 to Fig. 7

Supplementary Table 1 to Table 8

