## Supplemental Figure, Table, Information for "Mechanisms of perisynaptic astrocyte depolarization in response to neuronal activity"

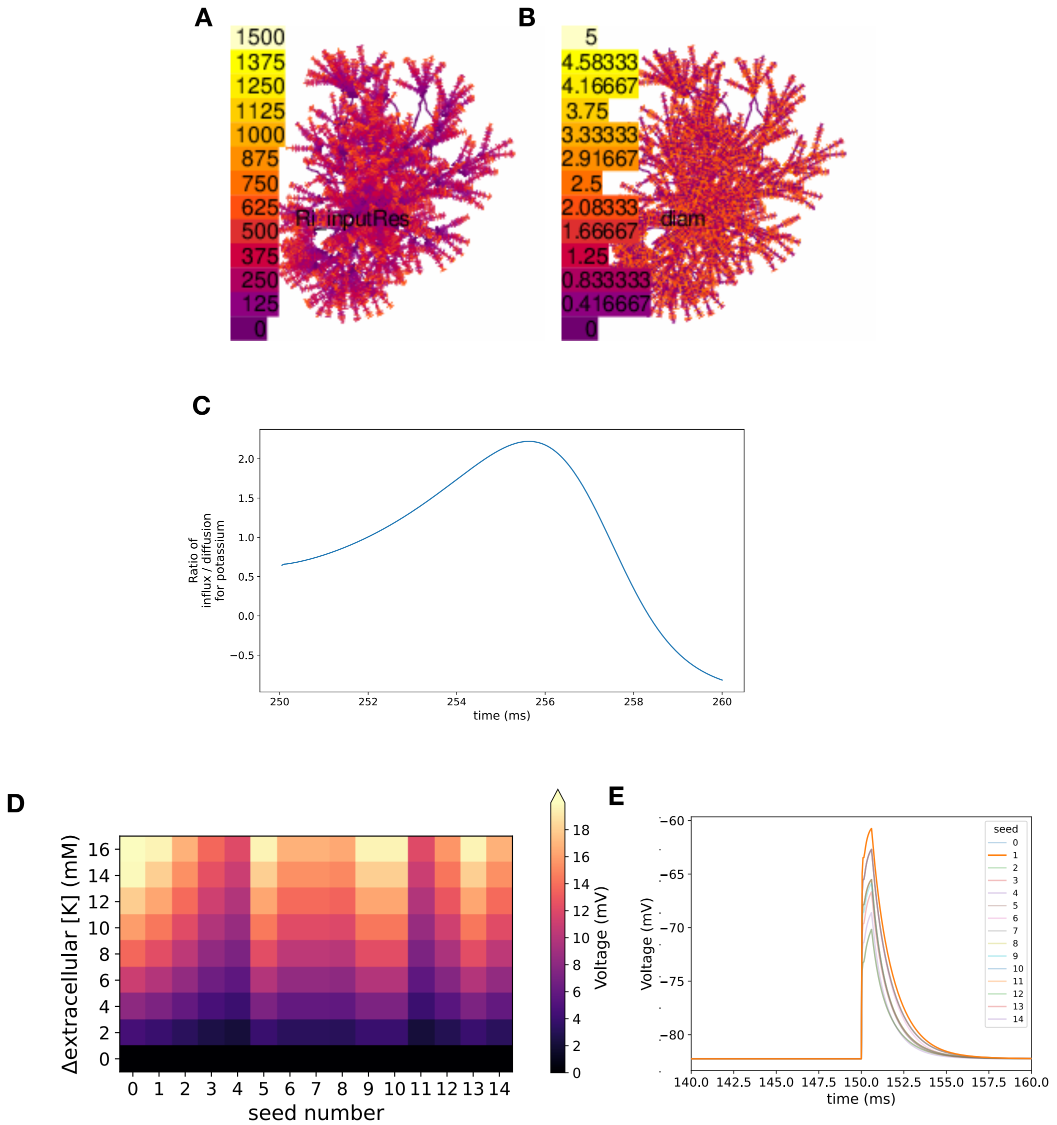

**Supplementary Fig. 1** Basic characteristics of the astrocyte whole-cell model

**Supplementary Fig. 1** Basic characteristics of the astrocyte whole-cell model
**(A,B)** Morphological characteristics of the astrocyte model. **(A)** Input resistance (color scale 0∼1500 MΩ) **(B)** diameter (color scale 0∼5 µm) of the hippocampal astrocyte morphology plotted for each section.
**C** Ratio of potassium clearance from the extracellular space. The ratio is between the astrocyte influx via Kir 4.1 and diffusion away from the astrocyte. The ratio plot shows the astrocyte response to 43 mM extracellular potassium increase with no glutamate. Potassium is increased at 150 ms for 100 ms and the ratio is recorded afterward. The ratio becomes greater than 1 soon after the extracellular potassium is applied, implying the astrocytic uptake is stronger. After the decrease of potassium, the efflux and diffusion away from the astrocyte equilibrate, resulting in a negative ratio. **(D)** Variability of PAP responses to local extracellular potassium changes dependent on location. Each location was selected randomly using different seeds. Maximum depolarization is indicated by the colors on the color bar, with various extracellular potassium changes. No depolarization of 20 mV can be achieved by 10 mM, and all PAPs require more potassium than estimated by the Nernst reversal potential. **(E)** Individual plots for PAP voltage response to 16 mM extracellular potassium changes. Each trace corresponds to the seed listed in the legend.

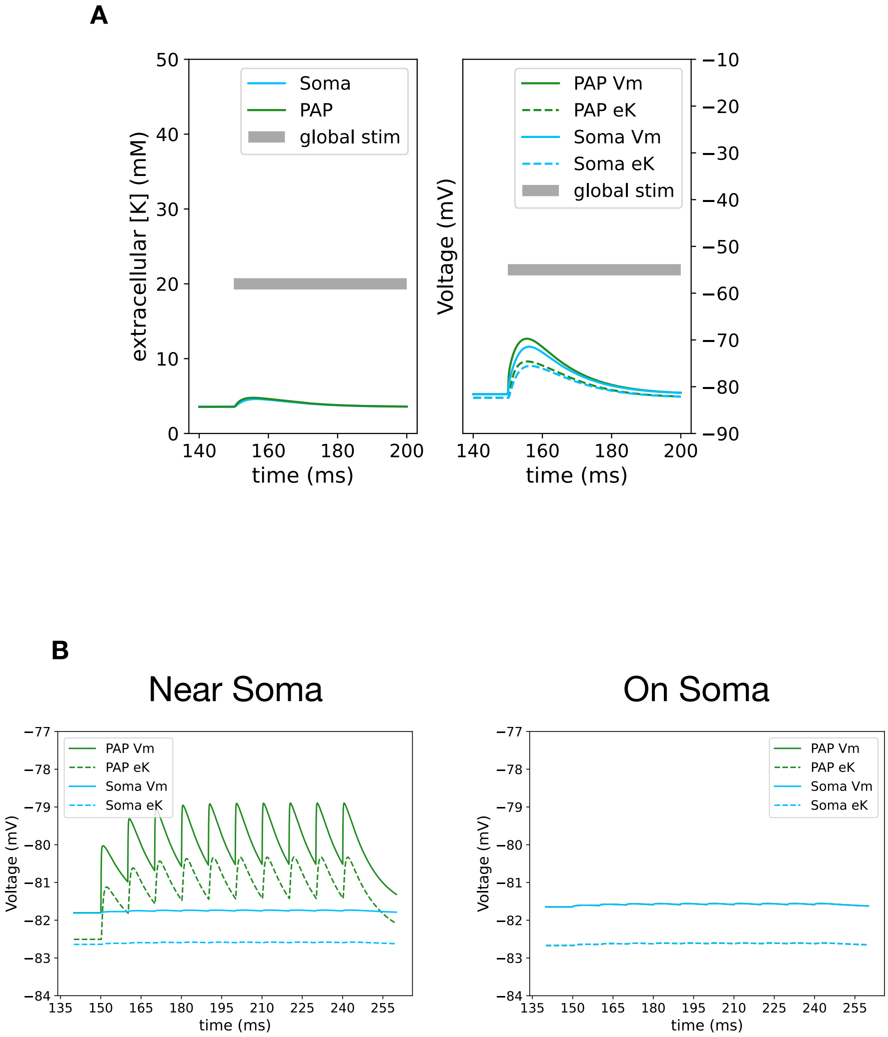


**Supplementary Fig. 2** GABA response to astrocyte in bath, near soma, and on soma application

**Supplementary Fig. 2** GABA response to astrocyte in bath, near soma, and on soma application
**(A)** Changes in extracellular potassium and membrane potential for both soma and PAP during the 1 mM GABA bath application over time. 1 $GABA_AR$ is placed per section of the model. **(B)** Membrane potential response in soma and PAP when PAP location is near soma (< 2 μm) or on soma. The stimulation protocol was that of 100 Hz for 10 pulses of sAP using GABA.

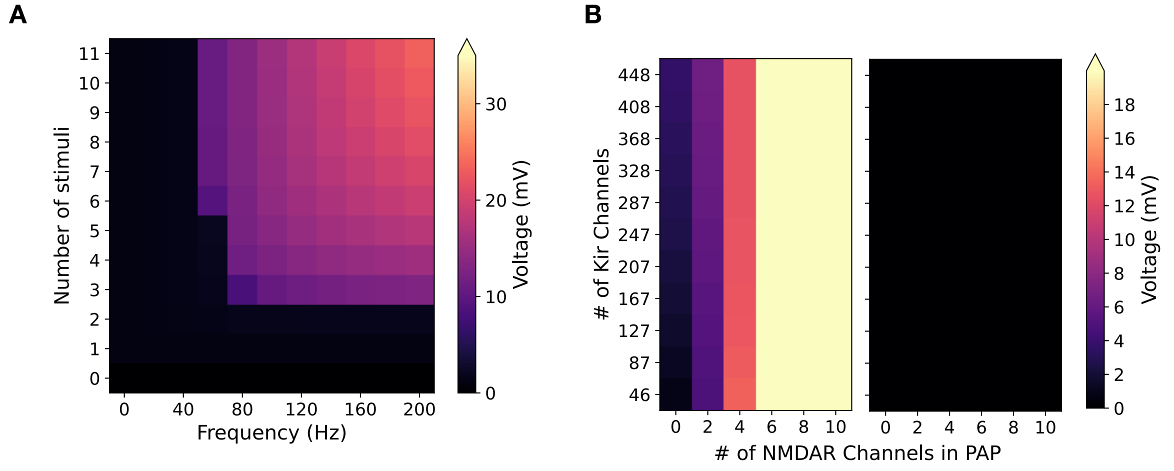


**Supplementary Fig. 3** Response of the model to various scenarios of the NMDAR stimuli

**Supplementary Fig. 3** Response of the model to various scenarios of the NMDAR stimuli
**(A)** Maximum depolarization achieved by various stimulation protocols. With larger Hz, larger astrocyte depolarization is achieved, which is also proportional to the number of stimuli. Protocols used throughout the paper were 10 pulses at 100 Hz. **(B)** Responses to combined stimulation of 10 PAPs. Left) Heat maps showing maximum depolarization for each count of the Kir 4.1 channel and NMDAR channel on the PAP for one of the 10 PAPs. Right) Heat maps showing maximum depolarization recorded at the soma for each count of the Kir 4.1 Channel and NMDAR channel on the PAP when 10 simultaneous PAP activations occur. There is no effect of simultaneous stimulation at the soma.

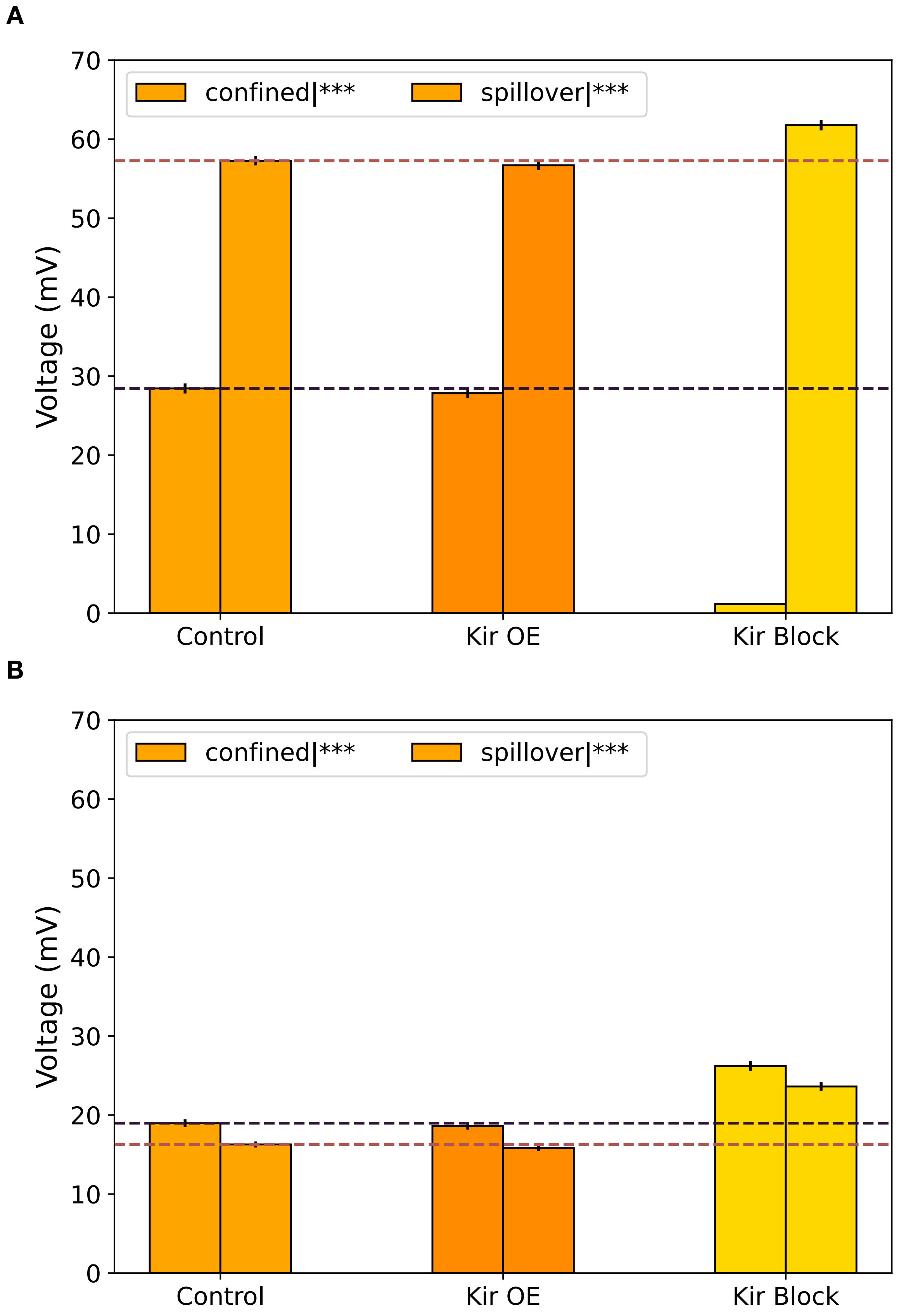


**Supplementary Fig. 4** Effect of Kir channel Over-expression/block on spillover effect for PAP depolarization

**Supplementary Fig. 4** Effect of Kir channel Over-expression/block on spillover effect for PAP depolarization **(A, B)** Comparison of peak depolarization with spillover over various Kir conditions. Kir 4.1 was either 482 channels at control, 873 at overexpression (OE), and 0 at Kir block (KO). Confined PAP lengths were defined as 0.3 µm while spillover was at 5 µm. (One-way ANOVA; ***P¡0.001) **(A)** Plots of peak depolarization for control and spillover with glutamate stimuli. OE significantly reduces peak depolarization while KO caused a significant increase in values for both conditions. **(B)** Plots of peak depolarization for control and spillover with GABA stimuli. OE does not significantly alter peak depolarization when compared to control for each respective condition. On the other hand, KO causes significant differences in peak depolarization. All spillover conditions significantly decrease when compared to confined conditions.

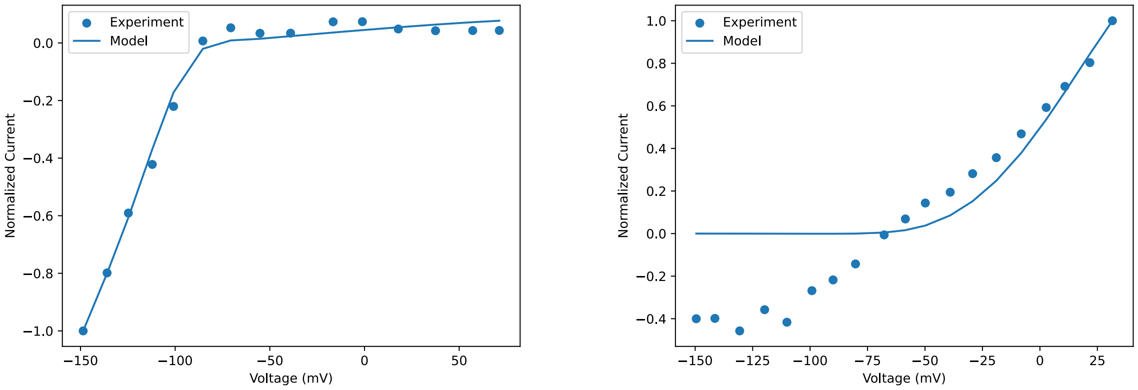


Kir 4.1 IV-curve TREK1 IV-curve

**Supplementary Fig. 5** Comparison of experimental data to model
Plots showing the normalized current-voltage relationship for the Kir 4.1 and TREK1. Each channel was fit to experimental results, where Kir 4.1 fits to a strong rectifying Kir 4.1 channel [61, 62] and TREK1 fits to experimental data used in Janjic et al. [51, 63]. The TREK1 model ignores inward current recordings following Janjic et al. [51]. All experimental conditions, such as intra/extracellular potassium concentrations and temperature, are considered in the model. For each plot, the x-axis is the holding membrane potential, and the y-axis is the peak inward current normalized to values recorded at the lowest holding potential, except for the TREK1 channel, which was normalized to the highest holding potential due to its nature. Note that goodness of fit was only considered for the relevant voltage range of the model.

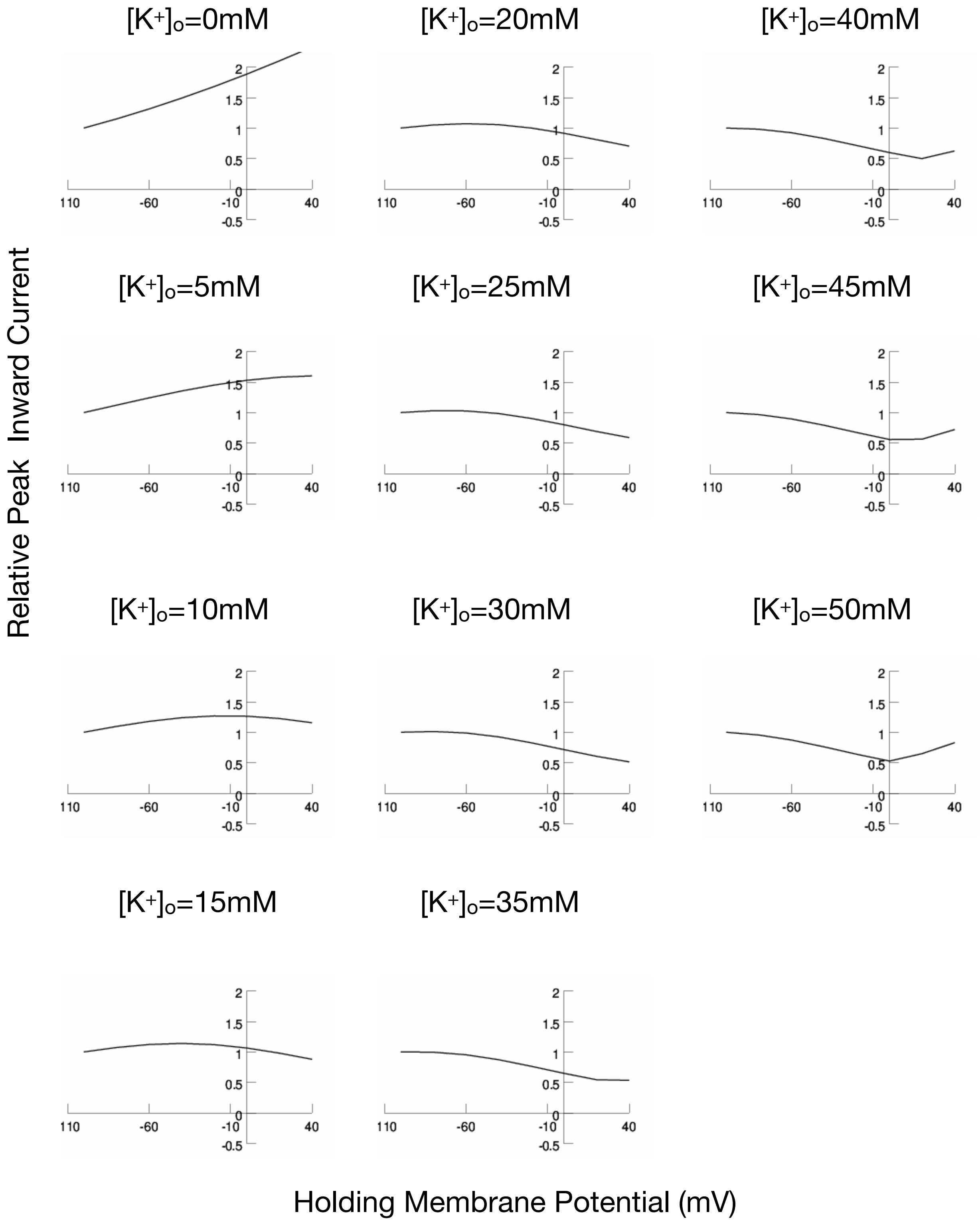


**Supplementary Fig. 6** GluT current dependence on extracellular potassium and holding voltage

**Supplementary Fig. 6** GluT current dependence on extracellular potassium and holding voltage
Plots showing the normalized current-voltage relationship for the GluT channel in our model. For each plot, the x-axis is the holding membrane potential, and the y-axis is the peak inward current normalized to values recorded at -110 mV. The peak inward current is proportional to the maximum amount of glutamate the astrocyte can uptake.

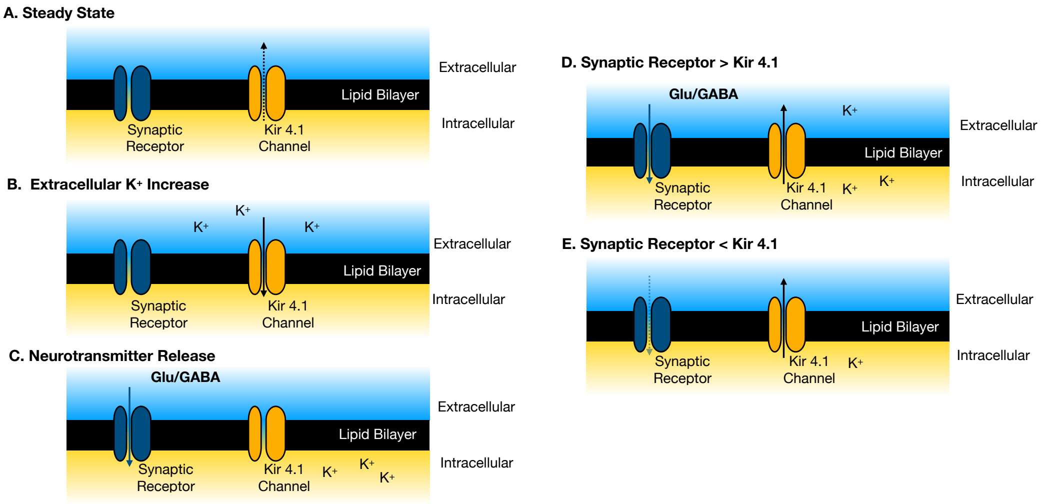


**Supplementary Fig. 7** Reaction scheme for synaptic receptor mediated depolarization

**Supplementary Fig. 7** Reaction scheme for synaptic receptor mediated depolarization
The cartoon depicts the direction of currents for Kir and synaptic receptors throughout the initial depolarization. All synaptic receptor currents are dark blue, and potassium currents are colored black. Dashed lines represent smaller currents. The cartoon depicts 5 stages that contribute to depolarization and repolarization. **(A)** Initial steady state of Kir 4.1 and synaptic receptor. Kir 4.1 exhibits a weak outward current maintaining RMP and extracellular potassium concentrations. **(B)** After application of extracellular potassium. As the reversal potential alters for Kir 4.1 upon increasing extracellular potassium, Kir 4.1 produces an inward current resulting in slight depolarization of the membrane. **(C)** Depolarization driven by application of neurotransmitters. Neurotransmitters activate respective synaptic receptors, which cause large inward currents for both inhibitory and excitatory synapses. **(D)** Continued depolarization by the synaptic receptor causes potassium efflux. While the synaptic receptor currents are larger than the Kir 4.1 repolarization effect, potassium continues to accumulate in the extracellular space. **(E)** Repolarization of astrocyte membrane potential by Kir 4.1. As the synaptic receptor closes, the Kir 4.1 channel repolarizes the astrocyte to basal RMP conditions.

**Table 1**: Constants for whole-cell astrocyte model

| Parameter | Value | Unit | Note | Reference |
|---|---|---|---|---|
| Ra | 100 | $\Omega\,\mathrm{cm}$ | | [18] |
| cm | 0.8 | $\mu\mathrm{F\,cm^{-2}}$ | | [18] |
| $g_{passive}$ | $8.97 \times 10^{-5}$ | $\mathrm{S\,cm^{-2}}$ | | [18] |
| $e_{passive}$ | -85 | mV | | [18] |
| T | 307 | K | Matching experimental setup | [5] |
| R | 8.3145 | $\mathrm{J\,K^{-1}}$ | | |
| F | 96485.3321 | $\mathrm{C\,mol^{-1}}$ | | |
| dt | 0.01 | ms | | |
| $[K^+]_i$ | 80 | mM | set for potassium reversal potential of -91.7 mV before equilibrium phase | [64] |
| $[K^+]_o$ | initial: 3 post-equilibrium: 3.5 | mM | baseline;altered for experiments | [5] |
| $[Na^+]_i$ | 15 | mM | | [65] |
| $[Na^+]_o$ | 140 | mM | | [18] |
| $[Cl^-]_i$ | 30 | mM | | [56] |
| $[Cl^-]_o$ | 130 | mM | | [57] |
| $[Mg^+]_o$ | 1 | mM | | [66] |
| $[Glu^+]_i$ | 0.3 | $\mathrm{mM\,L^{-1}}$ | | [18] |
| $[Glu^+]_o$ | 20e-6 | $\mathrm{mM\,L^{-1}}$ | | [18] |
| $g_{Kir4.1}$ | 50 | pS | single channel conductance for $[K^+]_o = 40$mM | [49] |
| $g_{NMDAR}$ | 33 | pS | single channel conductance | [67] |
| K2P-TREK1 base permeability | $1.24{\times}10^{-8}$ | $\mathrm{cm^3\,s^{-1}}$ | | [51] |
| $g_{leak\_K}$ | $2.22{\times}10^{-8}$ | $\mu\mathrm{S\,\mu m^{-1}}$ | Janjic et al. leak conductance divided by surface area | [51, 60] |
| $g_{leak\_Na}$ | $8.85{\times}10^{-7}$ | $\mu\mathrm{S\,\mu m^{-1}}$ | fit for steady RMP and extracellular potassium | |
| $\tau_e$ | 3 | ms | timeconstant for extracellular clearance; fit to maintain physiological extracellular potassium concentration (2∼4 mM) | |
| $r_e$ | 100 | Å | effective thickness; chosen to match the synaptic cleft volume for a 0.6 µm hippocampal spine head diameter | [68] |

**Table 2**: Table for parameters of Kir 4.1 Model

| Parameter | Units | Description | Value | Comment |
|---|---|---|---|---|
| $g_{\text{Kir0}}$ | pS | Maximum conductance for Kir | 4.7 | calibrated to match 50 pS at extracellular potassium concentration of 40 mM [49] |
| $v_{halfl}$ | mV | Voltage at half of maximum $l$ | -98.92 | Fit to data in Stegen et al. [69] |
| $k_l$ | mV | Sensitivity of $l$ | 10.89 | Same as above [69] |
| $bt$ | $\text{ms}^{-1}$ | Control parameter for $\tau_l$ | $8.2\times10^{-2}$ | Fit to data in Janjic et al. [51] |
| $at$ | $\text{ms}^{-1}$ | Control parameter for $\tau_l$ | $6.1\times10^{-3}$ | Same as above [51] |

**Table 3**: Table for parameters of TREK-1 Model

| Parameter | Units | Description | Value | Comment |
|---|---|---|---|---|
| $\tau_{K2P}$ | ms | Activation time of K2P-TREK pore | 3.0 | from experimental data [63] |
| $v_s$ | mV | Slope factor, RT/F | 25.7 | At T = 298K, constant in whole study |
| $V^0_{12-K2P}$ | mV | Half-activation voltage $V_{12}$ at $[K^+]^b_o$ | -20.5 | $I_{K2P}$ model, $n_{K2P}$ activation |
| $S_{K2P}$ | | Scaling parameter in $n_{K2P}$ activation | 1.7 | Adjusted the shift of $n_{K2P}$ with [K+] |
| $[K^+]^b_o$ | mM | Baseline $[K^+]_o$ | 2.5 | |
| $P^B_{K2P}$ | $\mathrm{cm}^3\,\mathrm{s}^{-1}$ | Constant permeability of K+ in pores at baseline $[K^+]_o$ | 1.24e-08 | K+ permeability in NP formalism. Dimension reflects net currents instead of densities |
| k | | k-th power in K2P activation kinetics | 2 | HH-type formalism for $n_{K2P}$ |
| $z_{K2P}$ | | Charge valence for K2P pores, fixed | 1.0 | $I_{K2P}$ model and $n_{K2P}$ activation |

**Table 4**: Table of parameters for NMDAR model

| Parameter | Unit | Description | Value | Comment |
|---|---|---|---|---|
| $[Mg^{2+}]_O$ | mM | Extracellular magnesium concentration | 1 | From Spruston et al. [66] |
| s | mV | Voltage shift for magnesium block | -70 | Experimental parameter to recreate weak susceptibility to $Mg^+$ |
| IC50 | mM | Half maximal concentration | 500 | Experimental parameter to recreate weak susceptibility to $Mg^+$ |
| $\delta$ | | Relative electrical distance | 10 | Decreased to match low magnesium susceptibility |
| $E$ | mV | reversal potential | -0.7 | From Spruston et al. [66] |
| $k$ | $mV^{-1}$ | steepness of the $g_{VD}$-V graph | 0.007 | From Clarke et al. [70] |
| $v_0$ | mV | The $V_m$ at which $g_{VD,\infty}$ is zero | -100 | From Clarke et al. [70] |
| $w_B$ | | percentage of decay for B | 0.95 | |
| $w_C$ | | percentage of decay for C | $1 - w_B$ | |
| Time Constant | Unit | $Q_{10}$ | $T_0$ (°C) | References |
| $\tau_A$ | | $2.2 \pm 0.5$ | 31.5 | Hestrin et al. [71] |
| $\tau_B$ | | 3.68 | 35 | Korinek et al. [72] |
| $\tau_C$ | | 2.65 | 35 | Korinek et al. [72] |
| $\tau_g$ | | 1.52 | 26 | Kim et al. [73] |

**Table 5**: Table of parameters for $GABA_AR$ model

| Parameter | Unit | Description | Value | Comment |
|---|---|---|---|---|
| $V_{50}$ | mV | Voltage for half maximal rectification | -52 | From Schulz et al. [55] |
| $e_{Cl}$ | mV | chloride reversal potential | -40 | calculated from Nernst equation for model chloride conc. |
| $k$ | mV | steepness of the rectification factor | 3 | From Schulz et al. [55] |

**Table 6**: Time constants for $GABA_AR$ model

| Time Constant | Unit | value | References |
|---|---|---|---|
| $\tau_A$ | ms | .1 | From Schulz et al. [55] |
| $\tau_B$ | ms | 10 | From Schulz et al. [55] |

**Table 7**: Kinetic reaction rates for GluT

| Parameter | Unit | Description | Value | Comment |
|---|---|---|---|---|
| $k12$ | $\mathrm{L\,mM^{-1}\,ms^{-1}}$ | reaction rate from state 1 to 2 | 20 | Referenced from [74] |
| $k21$ | $\mathrm{ms^{-1}}$ | reaction rate from state 2 to 1 | 0.1 | Referenced from [74] |
| $k23$ | $\mathrm{L\,mM^{-1}\,ms^{-1}}$ | reaction rate from state 2 to 3 | 0.015 | Referenced from [74] |
| $k32$ | $\mathrm{ms^{-1}}$ | reaction rate from state 3 to 2 | 0.5 | Referenced from [74] |
| $k34$ | $\mathrm{ms^{-1}}$ | reaction rate from state 3 to 4 | 0.2 | Referenced from [74] |
| $k43$ | $\mathrm{ms^{-1}}$ | reaction rate from state 4 to 3 | 0.6 | Referenced from [74] |
| $k45$ | $\mathrm{ms^{-1}}$ | reaction rate from state 4 to 5 | 4 | Referenced from [74] |
| $k54$ | $\mathrm{L\,mM^{-1}\,ms^{-1}}$ | reaction rate from state 5 to 4 | 10 | Referenced from [74] |
| $k56$ | $\mathrm{ms^{-1}}$ | reaction rate from state 5 to 6 | 1 | Referenced from [74] |
| $k65$ | $\mathrm{L\,mM^{-1}\,ms^{-1}}$ | reaction rate from state 6 to 5 | 0.1 | Referenced from [74] |
| $k16$ | $\mathrm{L\,mM^{-1}\,ms^{-1}}$ | reaction rate from state 1 to 6 | 0.0016 | Referenced from [74] |
| $k61$ | $\mathrm{L\,mM^{-1}\,ms^{-1}}$ | reaction rate from state 6 to 1 | 2e-4 | Referenced from [74] |

**Table 8**: Paired t-test p-values between Confined and Spillover conditions

| Condition | p-value |
|---|---|
| Control (glutamate) | 2.0E-30 |
| NMDAR KO | 1.4E-15 |
| GluT KO (glutamate) | 1.2E-30 |
| NMDAR-GluT KO | 8.3E-16 |
| Control (glutamate) | 2.2E-17 |
| GABAR KO | 8.3E-17 |
| GluT KO (GABA) | 2.0E-17 |
| GABAR-GluT KO | 8.3E-16 |

### Supplementary Text

#### Supplemental text 1: Kir 4.1 channel model

The Kir 4.1 channel model has been based on Yim et al. [47]. It followed a conventional conductance-based model with a square-root dependence on extracellular potassium [48]. Conductance was defined as,

$$g_{\mathrm{Kir}} = g_{\mathrm{Kir0}}\sqrt{[K^+]_o} \tag{1}$$

where $g_{\mathrm{Kir0}}$ is the maximum conductance and $[K^+]_o$ is the extracellular potassium concentration. $g_{\mathrm{Kir}}$ was calibrated to fit experiments by Yang et al. [49], which were single channel conductance measured under 40 mM of extracellular potassium. The potassium concentrations in the astrocyte and the extracellular space gave the reversal potential ($ek$). Additionally, the model contained one Hodgkin-Huxley (HH) type activation particle $l$, which fits Kir 4.1 recordings used in Stegen et al. [69].

$$l = (l_{inf} - l)/\tau_l \tag{2}$$

Each $l_{inf}$ and $\tau_l$ are given by the equations below.

$$linf = 1/(1 + exp((v - v_{halfl})/k_l)) \tag{3}$$

$$\tau_l = 1/((at \cdot exp(-v/v_{halft}) + bt \cdot exp(v/v_{halft}))) \tag{4}$$

where $v_{halfl}, v_{halft}$ are the values of $l, t$ at the half of maximum, $at, bt$ are time constant control parameters fit to experimental data. The model is defined below, with all parameter values mentioned in the Supplementary Table 2.

$$i_{\mathrm{Kir}} = g_{\mathrm{Kir}} \cdot l \cdot (v - ek) \tag{5}$$

#### Supplemental text 2: Channel number estimates

Bovzic et al. reported the relative Kir4.1 immunolabeling fluorescent intensity, Kir 4.1 positive puncta size, and recorded currents in their paper, which was used to estimate the Kir 4.1 channel counts on the PAP [50].

Initially, they record a -1500 pA whole-cell current for Kir 4.1 response when astrocytes are voltage clamped at -90 mV (eK was -72 mV). Considering that a single channel conductance was 7.5 pS (calculated at 2.5 mM of extracellular potassium), we calculated that a total $1.111 \times 10^4$ channels are required to achieve these dynamics.

In Bovzic et al. the average surface area of Kir 4.1 expression is $30.041 \pm 0.001$ µm$^2$ [50]. This equals a channel density of $370e8 \pm 1e8$ channels cm$^{-2}$.

#### Supplemental text 3: K2P-TREK1 channel model

The reaction dynamics of TREK-1 were defined as follows, derived from the paper by Janjic et al. [51]. The only alteration was to change the whole-cell current (nA) to a current density (mA cm$^{-2}$), by dividing the whole-cell current by the estimated astrocyte surface area [60]. The authors incorporate a time constant to the gating particle $n$ by using a time constant $\tau_{K2P}$.

$$I_{K2P} = n^k I_{K2p-GHK} \tag{6}$$

$$= n^k P_{K2P} \frac{F^2 z_{K2P}^2}{RT} V_m \frac{([K^+]_i - [K^+]_o \exp(-z_{K2P}V_m/v_s))}{(1 - \exp(-z_{K2P}V_m/v_s))} \tag{7}$$

$$\frac{dn}{dt} = \frac{n_{K2P}(V_m) - n}{\tau_{K2P}} \tag{8}$$

$$P_{K2P} = P_{K2P}^b \left(1 + 0.85 \log_{10}\left(\left[K^+\right]_o / \left[K^+\right]_o^b\right)\right) \tag{9}$$

$$n_{K2P}\left(V, \left[K^+\right]_o\right) = \frac{1 - \left[K^+\right]_o / \left[K^+\right]_i}{\left(1 + \exp\left(-z_{K2P}F\left(V_m - V_{12\_K2P}\right)/RT\right)\right)} \tag{10}$$

$$V_{12-K2P}\left(\left[K^+\right]_o\right) = V_{12-K2p}^0 - S_{K2P}v_s \ln\left(\left[K^+\right]_o / \left[K^+\right]_o^b\right) \tag{11}$$

$z, F, R, T$ are thermodynamic constants corresponding to the valence of ions, Faraday's constant, gas constant, and absolute temperature. $n_{K2P}$ is the voltage dependent gating particle for this equation and $P_{K2P}$ permeability constant.

$\left[K^+\right]_i, \left[K^+\right]_o$ represents inside and outside potassium concentrations. $V_m$ is the membrane voltage. All parameter constants can be found in Supplementary Table 3.

Supplemental text 4: NMDAR channel model

The reaction dynamics of the NMDAR were defined as follows, based on Moradi et al. [30].

$$I = (w_C C + w_B B - A) \cdot g \cdot Mg \cdot (v - E) \tag{12}$$

$$\frac{dA}{dt} = -\frac{A}{\tau_A} \tag{13}$$

$$\frac{dB}{dt} = -\frac{B}{\tau_B} \tag{14}$$

$$\frac{dC}{dt} = -\frac{C}{\tau_C} \tag{15}$$

The current of the NMDA channel was modeled as a combination of three ODEs $A, B, C$ with specific weights $w_B, w_C$ for each. Each ODE had a time constant $\tau_A, \tau_B, \tau_C$. $g$ and $E$ were the conductance and reversal potential, according to Ohmic Law.

The NMDA channel differs from neuronal channels due to its weak Magnesium block, which was modeled as below.

$$Mg = \frac{1}{1 + [Mg^{2+}]_O \cdot {K_0}^{-1} \cdot \exp(z \cdot \delta \cdot F \cdot (-v + s) \cdot R^{-1} \cdot T^{-1})} \tag{16}$$

Here, $[Mg^{2+}]_O$ was the extracellular concentration of Magnesium. $z, F, R, T$ are thermodynamic constants corresponding to the valence of ions, Faraday's constant, gas constant, and absolute temperature. $\delta$ is the relative electrical distance for the magnesium block. $s$ was the factor added to shift the magnesium block effect in order to match the weaker susceptibility of astrocytic NMDARs. $K_0$ is the $IC_{50}$ for extracellular magnesium at 0 mM. Since magnesium blocks in astrocytic NMDARs are weaker than neuronal ones, parameters related to Magnesium block were altered to fit experimental results [52].

$g$ is also a dynamic value defined based on ODEs $A, B, C$ and additional changes to time constant, $\tau_A, \tau_B, \tau_C$ providing a HH-type gating scheme.

$$g = 1 + g_{VD} \tag{17}$$

$$\frac{\partial g_{VD}}{\partial t} = (w_C C + w_B B) \cdot (g_{VD,\infty} - g_{VD})/\tau_g \tag{18}$$

$$g_{VD,\infty} = k \cdot (v - v_0) \tag{19}$$

$$\tau_A = q_{10} \cdot (\tau_{A,0} + a_A \cdot \exp(-\lambda_A v)) \tag{20}$$

$$\tau_B = q_{10} \cdot (\tau_{B,0} + a_B \cdot \exp(\lambda_B v)) \tag{21}$$

$$\tau_C = q_{10} \cdot (\tau_{C,0} + a_C \cdot \exp(-\lambda_C v)) \tag{22}$$

$$q_{10} = Q_{10}^{(T_0 - T)/10} \tag{23}$$

where, $g_{VD}, v_0, \tau_g$ were the voltage-dependent conductance, and time constant respectively. The equations were changed to qualitatively mimic the voltage dependence seen in Lalo et al. [52]. $g_{VD,\infty}$ was the final value of $g_{VD}$ when $t = \infty$. $v_0$ was the membrane potential when $g_{VD,\infty} = 0$, $\tau_g$ is the time constant for conductance change, $k$ defines the rate of change. $g_{VI}$ the voltage-independent component has been changed from 1 µS to 33 pS to match single channel recordings.

The dynamic time constants were defined as an exponentially rising constant with initial value $\tau_0$ and $a, \lambda$ obtained from experimental data fitting. $\lambda$ is a decay constant for each equation. These time constants were multiplied by $q_{10}$, which converts these constants to temperature-dependent ones. $T_0$ is the reference temperature for the given time constants measured experimentally. For desensitization, the original model utilized fast and slow dynamics. As repetitive stimulation protocols used in the current model were faster than those used in the experimental data, the fast desensitization factor was neglected. Slow desensitization occurred at 2500 ms time constant.

These equations summarize the three exponential model by Moradi et al. [30], where specific values of parameters are shown below in Supplementary Table 4.

Supplemental text 5: $GABA_A$R model

The $GABA_A$R model was a classic two-state synaptic conductance model adapted from Schulz et al. [55] with changes to the reversal potential to fit astrocytic conditions. Parameters for each reaction rate are listed in the Supplementary Table 5 and Supplementary Table 6. The $GABA_A$R were modeled by the two-state model equations below.

$$I = g_{max} * f(v) * (B - A) * (v - e_{Cl}) f(v) = 1 + \frac{-0.75}{1 + \exp(\frac{v - V_{50}}{k})} \frac{dA}{dt} = -\frac{A}{\tau_A} \quad (24)$$

$$\frac{dB}{dt} = -\frac{B}{\tau_B} \quad (25)$$

The current of the GABA$_A$R channel was modeled as a combination of two ODEs $A, B$. Each ODE had a time constant $\tau_A, \tau_B$. $f(v)$ was the voltage dependent rectification factor for GABA$_A$R. $g_{max}$ and $E_{Cl}$ were the unitary conductance and chloride reversal potential, according to Ohmic Law. The main difference between the original model and our model was the change in reversal potential and unitary conductance, in order to match astrocyte dynamics.

<u>Supplemental text 6: Glutamate Transporter</u>

The glutamate transporter model was modified from Savtchenko et al. [18], with slight alterations such as changing the MODL file from a membrane mechanism to a point process by adding a synaptic weight and calculating the membrane surface area within the MODL file. The model was a 6-state kinetic model with the reactions listed below. Parameters for each reaction rate are listed in the Supplementary Table 7.

| Reaction | Forward reaction rate | Backward reaction rate |
|---|---|---|
| $C1 \leftrightarrow C2$ | $[Glu]_o \cdot k12 \cdot u(v, -0.1)$ | $k21$ |
| $C2 \leftrightarrow C3$ | $[Na]_o \cdot k23 \cdot u(v, 0.5)$ | $k32$ |
| $C3 \leftrightarrow C4$ | $k34 \cdot u(v, 0.4)$ | $k43$ |
| $C4 \leftrightarrow C5$ | $k45$ | $k54 \cdot [Glu]_i$ |
| $C5 \leftrightarrow C6$ | $k56 \cdot u(v, 0.6)$ | $k65 \cdot [Na]_i$ |
| $C6 \leftrightarrow C1$ | $Kin \cdot k61$ | $k16 \cdot u(v, 0.6) \cdot [K]_o$ |

The function $u(v, k)$ was defined as below.

$$u = exp(k \cdot v/(2 \cdot 26.7)) \quad (26)$$

where $v, k$ were dependent on the aforementioned reactions. Currents were calculated based on charges assigned to each reaction. Reaction between state $C1$ and $C6$ was designated for potassium current ik. Therefore, the model was tweaked so that these changes would affect the total potassium current in the NEURON model.

<u>Supplemental text 7: leak channels</u>

All leak channels followed conventional conductance-based equations.

$$I_i = g(V - e_i) \quad (27)$$

where $i$ denotes the type of ion (potassium, chloride, sodium) and $e$ was the specific reversal potential calculated from Nernst potential for a specific ion $i$. $g$ is the conductance, $V$ was the membrane potential and $I$ was the current of the leak channel. Parameters of conductance and reversal potential can be found in Supplementary Table 1.
